# Speech Sound Discrimination by Mongolian Gerbils

**DOI:** 10.1101/2021.12.10.471947

**Authors:** Carolin Jüchter, Rainer Beutelmann, Georg Martin Klump

## Abstract

The present study establishes the Mongolian gerbil (*Meriones unguiculatus*) as a model for investigating the perception of human speech sounds. We report data on the discrimination of logatomes (CVCs_1_ - consonant-vowel-consonant combinations with outer consonants /b/, /d/, /s/ and /t/ and central vowels /a/, /a:/, /ε/, /e:/, /I/, /i:/, /ɔ/, /o:/, /℧/ and /u:/, VCVs_2_ - vowel-consonant-vowel combinations with outer vowels /a/, /I/ and /℧/ and central consonants /b/, /d/, /f/, /g/, /k/, /l/, /m/, /n/, /p/, /s/, /t/ and /v/) by young gerbils. Four young gerbils were trained to perform an oddball target detection paradigm in which they were required to discriminate a deviant CVC or VCV in a sequence of CVC or VCV standards, respectively. The experiments were performed with an ICRA-1 noise masker with speech-like spectral properties, and logatomes of multiple speakers were presented at various signal-to-noise ratios. Response latencies were measured to generate perceptual maps employing multidimensional scaling, which visualize the gerbils’ internal representations of the sounds. The dimensions of the perceptual maps were correlated to multiple phonetic features of the speech sounds for evaluating which features of vowels and consonants are most important for the discrimination. The perceptual representation of vowels and consonants in gerbils was similar to that of humans, although gerbils needed higher signal-to-noise ratios for the discrimination of speech sounds than humans. The gerbils’ discrimination of vowels depended on differences in the frequencies of the first and second formant determined by tongue height and position. Consonants were discriminated based on differences in combinations of their articulatory features. The similarities in the perception of logatomes by gerbils and humans renders the gerbil a suitable model for human speech sound discrimination.

**Highlights:**

- Perceptual maps of vowels and consonants in Mongolian gerbils are derived
- Gerbils perceive vowels and consonants in the same way as humans
- Gerbils discriminate vowels based on frequency differences of the formants
- Gerbils discriminate consonants based on differences in articulatory features

## 1. Introduction

Accurate discrimination of speech sounds is essential for daily life communication. In order to investigate the processing and recognition of speech sounds, animals are often used as model organisms. In this context, Mongolian gerbils (*Meriones unguiculatus)* have proven to be suitable subjects for studies on speech sound discrimination and age-related changes in physiological functions such as the degeneration of speech processing (Eipert and Klump, 2020a, 2020b; Schebesch et al., 2010; Sinnott et al., 1997; Sinnott and Mosqueda, 2003; Sinnott and Mosteller, 2001). However, until now only small sets of speech sounds have been investigated in gerbils. In the present study, we tested a large set of speech sounds in gerbils and investigated their discrimination ability for different vowels and consonants in detail. The aim of this study was to investigate young normal-hearing Mongolian gerbils as an animal model in order to obtain a reference data set regarding the processing of speech sounds. In combination with further studies in old gerbils, this knowledge may contribute to the understanding of speech processing in human listeners having normal hearing or showing different degrees of hearing loss (Dubno et al., 2008; Füllgrabe et al., 2015; 2007; Humes, 1996; van Rooij and Plomp, 1990).

### 1.1. Speech Sounds

Human speech comprises different types of speech sounds, of which vowels, consonants and logatomes are most important for the present study. Vowels are essential elements of speech sounds and they exhibit a harmonic structure (Hillenbrand and Nearey, 1999; Klatt, 1982; Nearey, 1989). Further, they are represented by complex spectral patterns that consist of a series of so-called formants, which are emphasized harmonics due to resonances in the vocal tract that can range from approximately 250 Hz to 3 kHz (Sinnott et al., 1997). The constellation of formants characterizes the specific vowel and its spectral shape (Schebesch et al., 2010). Human vowels are primarily discriminated based on the frequency relationship of the first formant (F1) and the second formant (F2) of the vowel spectrum (Assmann et al., 1982; Chistovich, 1985; Delattre et al., 1952; Liberman et al., 1967; Ohl and Scheich, 1997; Peterson and Barney, 1952). Yet, the formant frequencies of the same vowel can be different for different speakers, depending on the length of the individual’s vocal tract (Schebesch et al., 2010). Furthermore, vowels can be classified according to their articulatory configurations that range from open to close for the manner of articulation on the one hand and from front to back for the place of the articulation on the other hand (Ladefoged and Johnson, 2011). In contrast to vowels, consonants are characterized by a closure at one or more points in the vocal tract. They are typically classified on the basis of the three properties voicing, manner of articulation, and place of articulation (Ladefoged and Johnson, 2011). The 12 consonants that were used in the present study can be classified as voiced or unvoiced for the articulatory feature voicing, while the manner of articulation can be either plosive, nasal, fricative, or lateral approximant. The place of articulation ranges from labial to coronal to dorsal (International Phonetic Association, 2015). Logatomes are a higher-level type of speech sounds than vowels and consonants. Logatomes are nonsense syllables that consist of multiple vowels and consonants. They are commonly used in acoustic experiments and can provide a precise differentiation of phonemic confusions (Welge-Lüssen et al., 1997). Since neither gerbils nor humans associate any meaning with logatomes, these are particularly suitable for the comparison of speech sound discrimination between humans and animals.

### 1.2. Psychoacoustic Experiments

Psychoacoustical experiments offer the opportunity to learn more about the discriminability of different speech sounds without involving invasive methods. Instead, behavioral responses associated with different acoustic stimuli are used to draw conclusions about the internal representations and processing mechanisms of speech. Mongolian gerbils (*Meriones unguiculatus*) show a good low-frequency hearing with a similar sensitivity as humans for frequencies between 1 and 4 kHz (Ryan, 1976), which makes them suitable as an excellent animal model in hearing research and especially for the examination of human speech sounds. Besides their suitability as a model in hearing research, gerbils are also suitable subjects for studying the effects of aging on physiological functions, because they only live three to four years (Cheal, 1986). Moreover, old gerbils show increased auditory thresholds, age-related deficits in the representation of temporal structures of sounds and characteristics of strial and neural presbycusis (Gleich et al., 2016; Hamann et al., 2002; Heeringa et al., 2020; Hellstrom and Schmiedt, 1990; Kessler et al., 2020; Laumen et al., 2016; Tarnowski et al., 1991). Consequently, they have been used to study age-related hearing loss and its different structural and functional aspects (Gleich and Strutz, 2012). Another advantage of the usage of gerbils as an animal model is that they can easily be trained to respond to acoustic stimuli using operant conditioning (Ryan, 1976; Sinnott, 1995; Tolnai et al., 2017). Furthermore, the neural representation of speech sounds in animals is sufficiently rich for research on the processing of complex auditory stimuli (Mesgarani et al., 2008). In line with this, it has been shown that rodents and other animals can be trained to discriminate phoneme pairs categorically and that they are able to generalize what they have learned to novel situations (Kuhl and Miller, 1975). Gerbils have proven their ability to classify vowels correctly despite the variability of different speakers (Schebesch et al., 2010). Altogether, gerbils are well suited as small-mammal models for the processing of spectral cues in human speech sounds (Sinnott and Mosteller, 2001) and the categorization of these stimuli.

Here we address the following questions: (1) How do gerbils assess the variability in different speech sounds and how is the internal speech representation altered under different conditions? (2) Which stimulus features lead to salient differences between different vowels and consonants in gerbils? (3) How does speech sound discrimination differ between gerbils and humans?

## 2. Materials and Methods

### 2.1. Subjects

The experiments were conducted with four Mongolian gerbils (*Meriones unguiculatus*) that were bred and raised in the animal facilities of the University of Oldenburg and originated from animals obtained from Charles River laboratories. Over the time course of data collection, the animals were between 8 and 21 months old. Initially, at the age of 5 to 8 months, the gerbils were trained to discriminate pure tones and harmonic complexes in noise, before they were exposed to human speech sounds as stimuli. The gerbils were housed in pairs (two females and two males, respectively) and their cages contained litter and paper towels, cardboard and paper tubes, and wood sticks for nesting, hiding, and gnawing, respectively. All gerbils were trained five to six days a week. The gerbils were food-deprived for the period of the experimental data acquisition to increase the motivation during the experiments and only small amounts of rodent dry food were given outside of the experiment every evening. During the experimental sessions, the animals received custom-made 10-mg-pellets as rewards. The gerbils had unlimited access to water and their body weight and general condition were checked every day. The gerbils’ body weights during testing ranged from 63 to 87 g, corresponding to approximately 90% of their free feeding weights. All gerbils showed normal hearing abilities as assessed by auditory brainstem response measurements (ABRs) during behavioral testing. The care and treatment of the animals were approved by the Niedersächsisches Landesamt für Verbraucherschutz und Lebensmittelsicherheit (LAVES), Lower Saxony, Germany, permit AZ 33.19-42502-04-15/1990. All procedures were performed in compliance with the NIH Guide on Methods and Welfare Consideration in Behavioral Research with Animals (National Institute of Mental Health, 2002).

### 2.2. Experimental Setup

The experiments were conducted in two functionally equivalent setups. Both setups were situated in sound-attenuating chambers (IAC, Industrial Acoustics Company, 403-A or 401-A), with walls, ceiling, and floor covered with sound-absorbing foam (Pinta acoustic, PLANO 50/0 covered with PYRAMIDE 100/50 willtec). The reverberation times (T_60_) in the 403-A and 401-A chambers were 12 ms and 30 ms, respectively, suggesting nearly anechoic conditions. In the center of each setup was a custom-built elongated platform approximately one meter above the ground with a pedestal in the middle. A food bowl that was connected to an automatic custom-built feeder was positioned at the front edge of the platform. In front of the platform was a loudspeaker (Canton, Plus XS.2) driven by a power amplifier (Rotel, RMB-1512 or RMB-1506). The movements of the animal on the platform as well as its position on the pedestal were detected by two custom-built light barriers. The light barriers were connected via a signal processor (Tucker-Davis Technologies, RP2.1) to a Linux computer (Dell, OptiPlex 780 or OptiPlex 5040) with custom software that was used to monitor the experiments. A sound card (RME, Hammerfall DSP Multiface II) produced the signals, which had been pre-filtered offline with a 1024_th_ order FIR filter for a flat transfer function of the sound system. An infrared camera (Conrad Electronics, 150001 C-MOS) above the platform allowed visual control of the gerbil during testing. To calibrate the setup, the overall sound pressure level was measured two times a week for a reference frequency of 1 kHz. To do so, the microphone of a sound level meter (Brüel & Kjær, 2238 Mediator) was positioned approximately where the gerbil’s head was located during the experiments and, if necessary, the sound pressure level was adjusted with a resistance attenuator (TEXIO, RA-920A).

### 2.3. Stimuli

The logatomes that were used for the experiments were obtained from the *Oldenburg Logatome Corpus* (*OLLO*) (Meyer et al., 2010) and were spoken by two female and two male German speakers in normal speaking style and without a dialect. In total, the stimulus set for the present study comprised 40 CVCs (consonant-vowel-consonant combinations) and 36 VCVs (vowel-consonant-vowel combinations). The CVCs were combinations of the outer consonants /b/, /d/, /s/ and /t/ and the central vowels /a/, /a:/, /ε/, /e:/, /I/, /i:/, /ɔ/, /o:/, /℧/ and /u:/, with the outer phonemes being identical within one logatome. The VCVs consisted of the outer vowels /a/, /I/ and /℧/, combined with the central consonants /b/, /d/, /f/, /g/, /k/, /l/, /m/, /n/, /p/, /s/, /t/ and /v/. Only the discriminability between logatomes with the same outer phoneme were tested. A noise-masker with speech-like spectral properties (*International Collegium for Rehabilitative Audiology noise track 1* – *ICRA-1*) (Dreschler et al., 2001) was presented continuously during the sessions with 60 dB sound pressure level (SPL). The temporal and spectral properties of the *ICRA-1* noise closely resemble those of real-life communication, similar to a loud cocktail party noise (Meyer et al., 2010). Two different signal-to-noise ratios (SNRs), namely +5- and +15-dB SNR, were tested.

### 2.4. Procedure

The behavioral experiments employed a go/no-go paradigm. The animals were trained using operant conditioning with positive reinforcement (food pellets) to perform an oddball target detection task in which they were required to detect a change in a repeating background of sound. The reference logatome was repeated every 1.3 seconds and the speaker for each repetition was selected at random. After the animal positioned itself on the pedestal facing the loudspeaker to initiate a new trial, the next reference logatome after a random waiting time between one and seven seconds was replaced by a target logatome. The order of the trials with different target logatomes was randomized. The animal had to indicate a detected target logatome by jumping off the pedestal within 1.5 seconds from the onset of the target stimulus in order to achieve a hit and to get the food reward from the food bowl. To exclude trials in which the gerbils jumped off the pedestal by chance, the response time had to be at least 0.1 seconds. If the gerbil left the pedestal earlier, the animal was not rewarded, and the trial was repeated. Further, catch trials in which the reference stimulus was presented as a target stimulus were introduced to determine the false-alarm rate as a measure of spontaneous responding. The proportion of catch trials in the CVC- and VCV-conditions amounted to 30.8% and 26.7%, respectively. Eight CVC-conditions and six VCV-conditions were tested in order to investigate the influence of the different outer phonemes in combination with the two different SNRs on the discriminability of the logatomes. Each condition was divided into eight sessions with 65 and 90 trials for CVCs and VCVs, respectively. One session consisted of five blocks with 13 trials in CVC-conditions and six blocks with 15 trials in VCV-conditions, with changing reference-stimuli between the different blocks. Sessions could be stopped at any time and restarted later to be completed. Sessions typically lasted between 20 and 60 minutes and were conducted without visible light in the chamber. Response latencies and detection probabilities (hit rates) were measured for the different stimulus sets and conditions for all gerbils. In order to ensure that the gerbils were performing the experiments under stimulus control, values for the sensitivity-index *d’* were calculated, applying the inverse cumulative standard normal distribution function *Φ*^*−1*^ to the hit (*H*) and false-alarm rates (*FA*): *d’* = *Φ*^*−1*^(*H*) – *Φ*^*−1*^(*FA*) (Green and Swets, 1966). As a validity criterion, the averaged *d’* of a complete session had to be larger than 0.5 in order to prevent floor effects, otherwise the session had to be repeated. This criterion corresponds to maximum false-alarm rates of 30% and 21% for typically observed hit rates in CVC- and VCV-conditions, respectively. However, the d’ values were much larger for the vast majority of the sessions.

### 2.5. Human Data

In order to be able to draw conclusions about the comparability of speech sound discrimination in gerbils and humans, also human data on the discrimination of speech sounds was obtained. In a similar experiment with the same experimental paradigm, the discriminability of a subset of the same logatome stimuli was tested in five (4 female, 1 male) young adult, normal-hearing, German native speaking human subjects (age range 22-29 years) in the course of a student practical course (unpublished data). The experimental procedure for the humans was similar to that of the gerbils. However, stimuli were presented via headphones (Sennheiser, HDA 200) to the subjects seated in a sound-attenuating chamber (IAC, Industrial Acoustics Company, Mini 250) that responded using a touch screen (Elo, 1542L 15” Touchscreen Monitor). In the CVC-experiments for the human subjects, CVCs with the outer consonant /b/ and the same central vowels as in the experiments with the gerbils were presented at a SNR of -7 dB SPL. For the VCV-experiments, the same logatomes as in the VCV-conditions for the gerbils were used and presented at -7 dB SPL. The experiments were done with the understanding and written consent of each subject following the Code of Ethics of the World Medical Association (Declaration of Helsinki).

### 2.6. Data Analysis

#### 2.6.1. Confusion Matrices

The data analysis was conducted with *IBM SPSS Statistics 26* and *Microsoft Excel 2016*. Confusion matrices (CMs) served as the basis for the analysis. The scores in the cells of the CMs were ranks that were based on the response latencies of the subject, which quantify its discrimination performance for each combination of reference and target stimuli. The smaller a response latency rank, the faster the subject responded to the target stimulus. Thus, small response latency ranks indicate salient differences between reference and target stimuli, whereas large response latency ranks suggest a minor discriminability. CMs are based on all sessions of the corresponding condition for one individual. All latencies (per trial) of one condition and one individual were pooled for rank transformation. Missed responses were inserted as their maximal response time of 1.5 seconds and the average rank of all corresponding ranks was assigned to tied values. Subsequently, the ranks were averaged within the groups defined by background/target combination, resulting in a CM of ranks. With the help of CMs, it can be characterized how well a specific logatome was discriminated from other logatomes.

#### 2.6.2. Multidimensional Scaling

For evaluating the CMs, multidimensional scaling (MDS) was used. In animal psychophysics, MDS can provide insight into the unknown acoustic dimensions in the perception of non-human animals (Dooling and Okanoya, 1995) and the underlying neuronal processes of the discrimination among complex acoustic stimuli such as speech (Okanoya and Dooling, 1988). Furthermore, data from different subjects can be analyzed together using an individual difference model of scaling, offering the possibility to generate a common representation of the perceived stimuli for all subjects. In the present study, MDS was used to translate the differences in response latency ranks between stimuli to distances in multidimensional space in so-called perceptual maps. These perceptual maps represent the perceived stimulus similarity by spatial proximity of the logatomes (Dooling and Okanoya, 1995), such that short response latencies are reflected by large distances representing salient differences between stimuli, whereas long response latencies are reflected by short distances representing high similarity between stimuli. We used the MDS program *PROXSCAL* (Busing et al., 1997) with a generalized Euclidean scaling model. The perceptual maps for the present study were arranged in a two-dimensional space. The perceptual dimensions and the distances between vowels and consonants can be correlated to features of the speech sounds in order to interpret the perceptual maps. The *Dispersion Accounted For* (*DAF*) was used to evaluate the goodness of fit of the perceptual maps to the underlying data. The *DAF* provides a measure for the proportion of the sum of the squared disparities (transformed proximities) that is explained by the distances in the MDS solution (Borg et al., 2010).

#### 2.6.3. Hierarchical Cluster Analysis

Beyond the MDS, hierarchical cluster analyses were performed using the coordinates of the perceptual maps obtained from the MDS. Cluster analyses are used to describe the structure of similarity data and to cluster different stimuli into groups that should reveal meaningful features of the stimuli (Dooling and Okanoya, 1995). In contrast to MDS, which yields continuous dimensions (Dooling and Okanoya, 1995), cluster analyses produce discrete representations of the stimuli in a stimulus set. The clustering is based on the average linkage between the groups according to Euclidean distance metric. For this purpose, dendrograms were generated that illustrate the arrangement and dissimilarity of the clusters. The height of a dendrogram reflects the distance between the clusters and the dendrograms can be ‘cut’ at particular levels, resulting in specific numbers of clusters (Everitt and Skrondal, 2010). To ensure the comparability of the results of the different conditions, the same sectional planes at the dissimilarity levels 9 and 11 on the x-axis of the dendrograms were chosen for all CVC- and VCV-dendrograms, respectively. The resulting clusters were marked by black circles in all perceptual maps.

#### 2.6.4. Statistical Analysis of *d’*-Values, Response Latencies and Perceptual Distances

General data analysis and significance testing were conducted on the dependent variables *d’*-value and response latency using a two-way MANOVA to test for differences of means between the factors logatome type (CVCs vs. VCVs) and SNR (5 vs. 15 dB SNR). Response latencies and *d’*-values were averaged within conditions before entering into the MANOVA. Differences between means of perceptual distances derived from MDS were evaluated regarding the similarity of articulatory features of phoneme pairs (similar vs. different characteristics of different articulatory features) by unpaired two-sample t-tests. Further, one-way ANOVAs were used to test for significant differences between mean perceptual distances of phoneme pairs depending on the number of shared articulatory features. A Bonferroni correction was applied for post-hoc analysis to correct for multiple comparisons.

## 3. Results

Mongolian gerbils were trained to discriminate between different logatomes. We used CVCs and VCVs with varying central vowels and consonants, respectively. Generally, we found that the discrimination of CVCs was much easier for the gerbils than the discrimination of VCVs. A two-way MANOVA with logatome type (CVC, VCV) and SNR (+5 dB, +15 dB) as fixed factors and *d’*-value and response latency as dependent variables showed a significantly higher mean *d’*-value for the CVC-conditions in comparison to the VCV-conditions (*F*(1,55) = 42.677, *p* < 0.001; see Figure 1, panel A). Thus, a significantly higher proportion of vowel changes (54%) was detected by the gerbils in comparison to the proportion of detected consonant changes (38%). In line with this, the mean response latency (see Figure 1, panel B) was significantly shorter for the detection of changes between CVCs than between VCVs (*F*(1,55) = 55.607, *p* < 0.001). No significant change in the mean *d’* value could be assessed for the different SNRs (*F*(1,55) = 0.866, *p* = 0.356; see Figure 1, panel A). However, a significant difference in mean response latency was observed between the different SNRs (*F*(1,55) = 6.107, *p* = 0.017; see Figure 1, panel B) with higher SNRs associated with shorter response latencies. No significant interaction effect between logatome type and SNR on the dependent variables *d’* and response latency was observed (*F*(2,51) = 0.926, *p* = 0.403).

**Figure 1:**
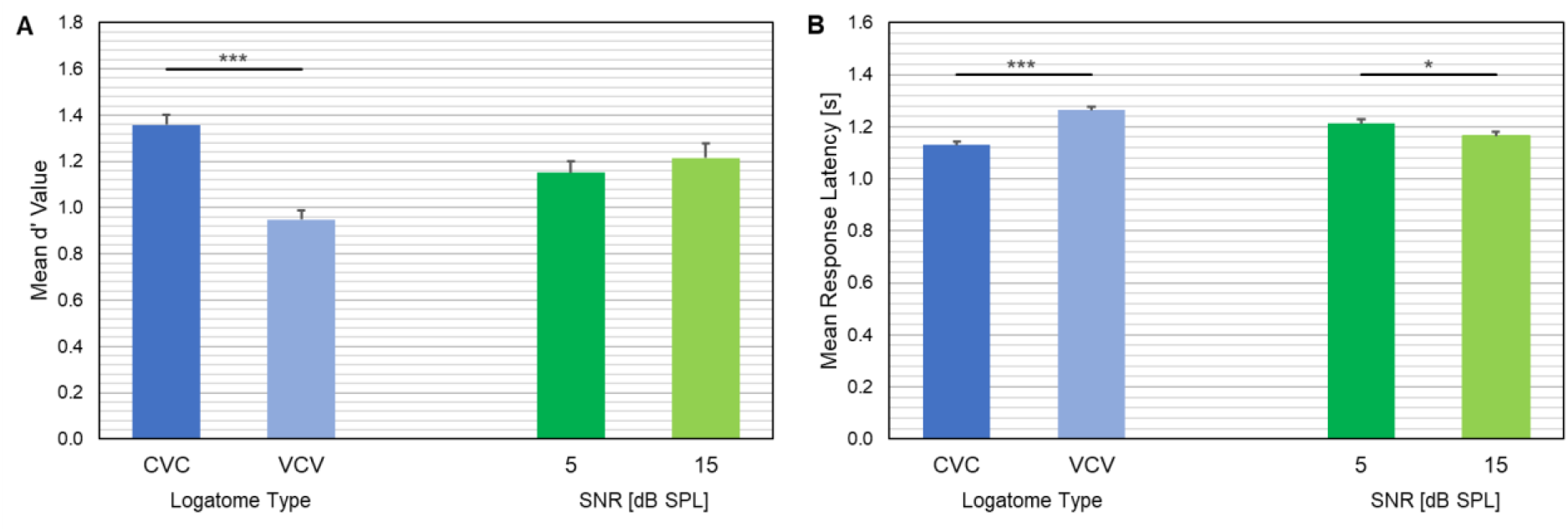
Mean d’ values (A) and mean response latencies (B) for the different logatome types and SNRs. Error bars indicate the standard error of the mean. Significant differences are marked by stars (*: p < 0.05, ***: p < 0.001).

### 3.1. Consonant-Vowel-Consonant Combinations

Perceptual maps reflecting the gerbils’ internal representations of the logatomes were generated. These were similar for the different CVC-conditions and individuals (see Supplementary Material A - Supplementary Material H). Further, the spatial arrangement of the vowels as well as the cluster constellations and the distances between the vowels were similar for all different outer consonants. Similarly, only minor differences were found between the perceptual maps for different SNRs in the CVC-conditions.

In the different perceptual maps of all CVC-conditions, the vowels /a/ and /a:/ made up one cluster in the hierarchical cluster analysis. The position of the other vowel clusters in relation to this cluster was quite similar for all CVC-conditions. There were always two (one time three) clusters with different constellations of the vowels /ε/, /e:/, /I/ and /i:/ positioned towards the negative range of dimension 1 and two to three clusters with different constellations of the vowels /ɔ/, /o:/, /℧/ and /u:/ positioned towards the positive range of dimension 1. For the vowels /ε/, /e:/, /I/ and /i:/, the distance between the cluster of /a/ and /a:/ and the individual vowels was smallest for /ε/, next smallest for /I/ and largest for /e:/ and /i:/ in all perceptual maps. For the vowel /ɔ/, /o:/, /℧/ and /u:/, the vowel /ɔ/ lay closest to the cluster of /a/ and /a:/ in seven out of eight conditions, whereas the other three vowels showed varying distances to the cluster of /a/ and /a:/. The vowel /u:/ was arranged furthest away from the cluster of /a/ and /a:/ in six out of eight conditions. The *DAF* values for the perceptual maps of the different CVC-conditions indicated that between 94.1% and 95.8% of the sums of the squared disparities were explained by the distances in the MDS solutions. Thus, the two-dimensional perceptual maps reflected the gerbils’ perceptions of the vowels very well.

Because of the consistency in the perceptual maps of the different CVC-conditions, a shared analysis for all CVC-conditions was carried out in order to create a generalized perceptual map of the ten tested vowels. The same cluster arrangements as described above for the different CVC-conditions were found for the shared perceptual map of all CVC-conditions and can be seen in Figure 2, panel A. The *DAF* value for the shared perceptual map of all CVC-conditions amounts to 93.9%.

**Figure 2:**
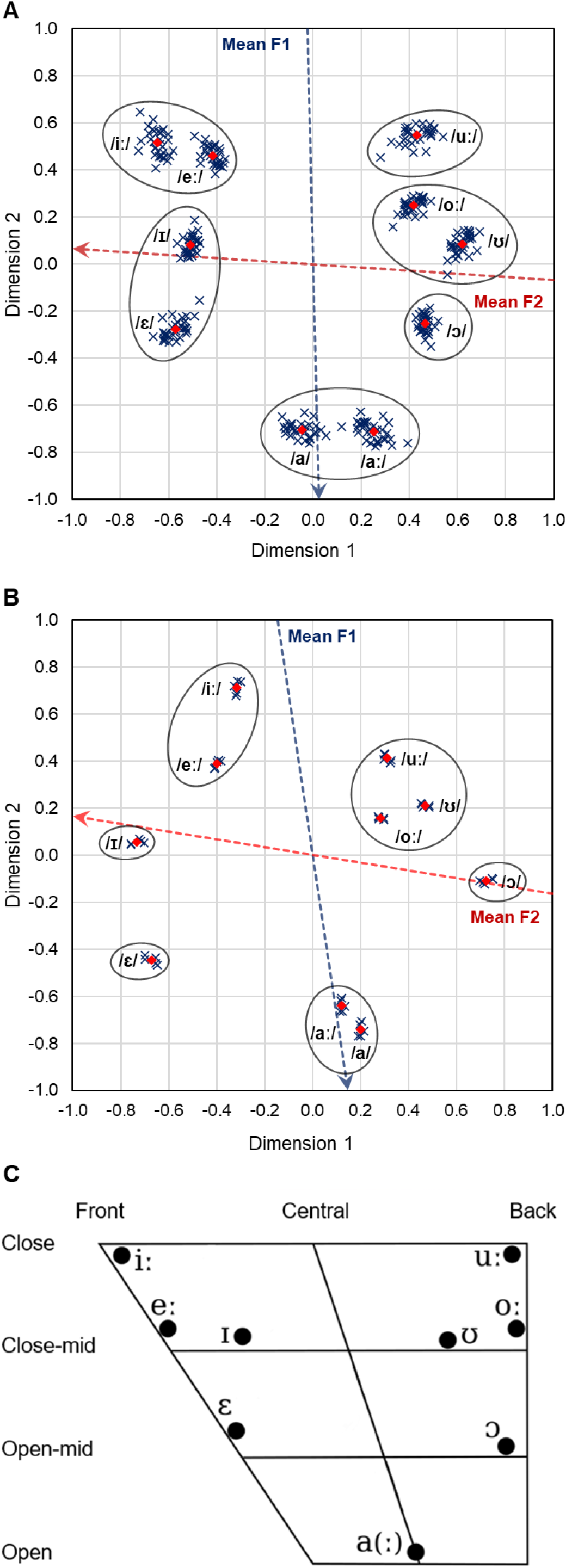
(A): Shared perceptual map of all CVC-conditions for four gerbils. Blue crosses: Individual coordinates for the different CVC-conditions. Red diamonds: Mean coordinates for all CVC-conditions. Black circles: Vowel clusters according to the hierarchical cluster analysis. Blue dotted arrow: Axis along which the mean F1 frequency of the vowels increases. Red dotted arrow: Axis along which the mean F2 frequency of the vowels increases. (B): Perceptual map of CVCs with the outer consonant /b/ at -7 dB SNR for five human subjects. Symbols and arrows as in (A). (C): Edited vowel chart for Northern Standard German (Kleiner et al., 2015) showing the articulatory configurations of the different vowels in the present study. The manner of articulation is determined by the tongue height (close: /i:/, /u:/ – close-mid: /e:/, /I/, /o:/, /℧/ – open-mid: /ε/, /ɔ/ – open: /a/, /a:/) and the tongue position regulates the place of articulation (front: /ε/, /e:/, /I/, /i:/ – central: /a/, /a:/ – back: /ɔ/, /o:/, /℧/, /u:/).

Apart from visualizing how the different vowels are internally represented in the gerbils, the perceptual map can further help to evaluate how the gerbils assessed the differences between the logatomes. As indicated by the dashed arrows in Figure 2, panel A, differences in spectral features – namely the frequencies of the formants – are important for the discrimination of vowels. We found that the frequency of F2 changed along dimension 1. In more detail, vowels with high dimension 1 values had low mean F2 frequencies and vowels with low dimension 1 values had high mean F2 frequencies. The mean F2 frequencies explained 89% of the variance in dimension 1 coordinates. A similar connection was found between the frequency of the first formant and dimension 2. Here, vowels that showed a high dimension 2 value had a low mean F1 frequency and vice versa. 96% of the variance in dimension 2 coordinates was explained by the mean F1 frequencies.

In addition to the spectral features of vowels, also their articulatory features are of interest for interpreting the perceptual maps of the CVC-conditions. If one compares the arrangement of the vowels in the shared perceptual map of all CVC-conditions with the arrangement of the vowels in the vowel trapezium for Northern Standard German (Figure 2, panel C), a high similarity can be observed. In the vowel trapezium, the vowels are represented according to their articulatory features with the place of articulation changing from left to right and the manner of articulation changing from the top to the bottom. The similar arrangement of the vowels in the vowel trapezium and the shared perceptual map of all CVC-conditions suggests that dimension 1, which is related to the frequency of the second formant, is determined by the place of articulation and dimension 2, which is related to the frequency of the first formant, is determined by the manner of articulation.

The spatial arrangement of the vowels in the perceptual map and the vowel clusters of the human subjects were similar to those of the gerbils (see Figure 2, panel B). This is further confirmed by the high correlation of the distances between the different vowel pairs in the perceptual maps of gerbils and humans, which is shown in Figure 3 (*R*_2_ = 0.71). As in the perceptual map of the gerbils, the frequencies of the first and second formant were significantly correlated with the second (*R*_2_ = 0.90) and first (*R*_2_ = 0.71) dimension of the humans’ perceptual map, respectively. Further, the same relation between spectral and articulatory features of vowels as in the perceptual map of the gerbils can also be found in the perceptual map of the human subjects. Accordingly, differences in the articulatory features of the vowels, which determine the formant frequencies, led to large distances in the perceptual maps of both gerbils and humans (see Figure 4). Unpaired two-sample t-tests for tongue position (same vs. different, see Figure 2C for classification of tongue position) and tongue height (same vs. different, see Figure 2C for classification of tongue height) revealed significantly larger mean perceptual distances for vowel pairs that are articulated with different tongue positions than for vowel pairs that share the same tongue position for gerbils (*p*_*Position*_ < 0.001; see Figure 4, panel A) and humans (*p*_*Position*_ < 0.001; see Figure 4, panel B). Different tongue heights did not lead to significant differences in perceptual distances for gerbils (*p*_*Height*_ = 0.296; see Figure 4, panel B) and humans (*p*_*Height*_ = 0.152; see Figure 4, panel A). However, this is only due to the large influence of the tongue position leading to long displacements between the vowels along dimension 1. The perceptual distances based only on the coordinates of dimension 2 were significantly smaller for vowel pairs that share the same tongue height than for vowel pairs with different tongue heights in gerbils (p < 0.001) and humans (p < 0.001). Moreover, one-way ANOVAs with the number of shared articulatory features (0, 1, 2) as fixed factor and the perceptual distance of the vowel pairs as dependent variable revealed that the more articulatory differences a vowel pair showed, the larger was the perceptual distance between the vowels for gerbils (*F*(2,44) = 18.426, *p* < 0.001; see Figure 4, panel C) and humans (*F*(2,44) = 16.741, *p* < 0.001; see Figure 4, panel D). Post hoc pairwise comparisons revealed significantly shorter perceptual distances for higher numbers of shared articulatory features for all comparisons in humans (*p*_0–1_ < 0.001, *p*_0–2_ < 0.001, *p*_1–2_ = 0.032; see Figure 4, panel D) and for all comparisons except for an increase from one to two shared articulatory features in gerbils (*p*_0–1_ < 0.001, *p*_0–2_ < 0.001, *p*_1–2_ = 0.077; see Figure 4, panel C).

**Figure 3:**
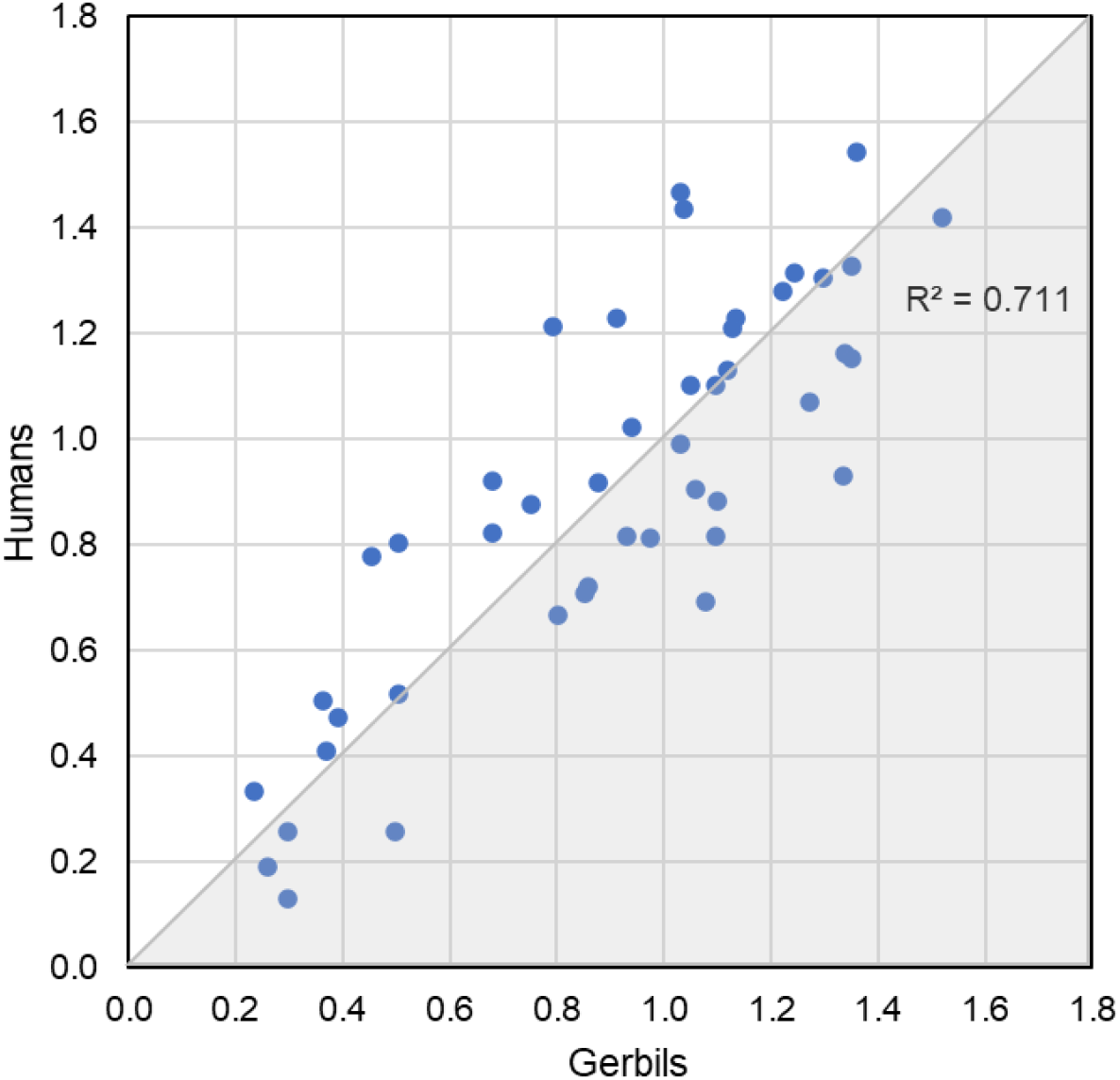
Perceptual distances of vowel pairs for humans vs. gerbils. Perceptual distances from humans are plotted against perceptual distances from gerbils for vowel pairs derived from MDS for CVC-conditions.

**Figure 4:**
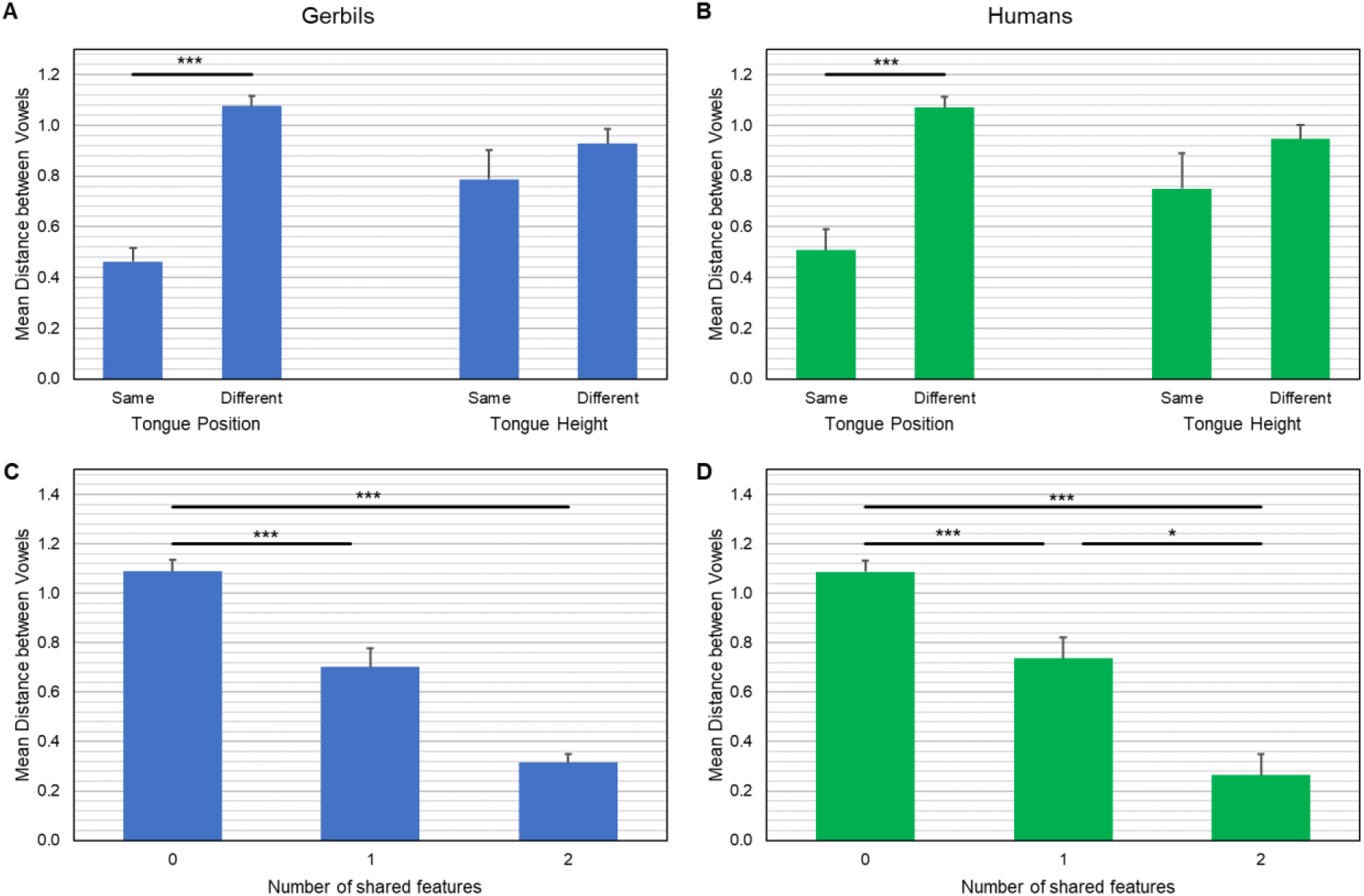
Mean perceptual distances of vowel pairs in relation to differences in tongue position and tongue height for gerbils (A) and humans (B). Mean perceptual distances of vowel pairs depending on the number of shared articulatory features for gerbils (C) and humans (D). Error bars indicate the standard error of the mean. Significant differences are marked by stars (*: p < 0.05, ***: p < 0.001).

### 3.2. Vowel-Consonant-Vowel Combinations

As for the CVCs, the spatial arrangements of the consonants of the VCVs in the perceptual maps were comparable for all individuals (see Supplementary Material I - Supplementary Material N). Moreover, the outer vowel had no large effect on the spatial arrangement of the perceptual maps of the different VCV-conditions. Similarly, only minor differences were found between the different SNRs in the VCV-conditions.

In the perceptual maps of all VCV-conditions, the central consonant /s/ made up a cluster alone in the hierarchical cluster analysis. The consonant /f/ also either made up a cluster alone (4 out of 6 conditions) or was clustered together with /v/, another labial fricative (2 out of 6 conditions). In 50% of the conditions, the central consonant /t/ also made up its own cluster, whereas it was clustered with different other consonants in the other 50% of the conditions. All other consonants were clustered together in different constellations with other consonants for the different conditions. In total for all six conditions, there were 24 clusters with more than one consonant. From these ‘multi-consonant clusters’, 96% of the clusters only contained consonants that shared a least one articulatory feature (voicing, manner of articulation or place of articulation) with every other consonant in the cluster. The consonants in 42% of the multi-consonant clusters even had two articulatory features in common with every other consonant of the cluster. Only one very large cluster that consisted of 5 consonants for the VCV-condition with the outer vowel /℧/ at +5 dB SNR (see Supplementary Material M) did not show common features among all involved consonants. Furthermore, clusters never comprised more than two different characteristics of one articulatory feature (e.g. not more than two different manners of articulation within one cluster). According to the *DAF* values, between 89.8% and 92.5% of the sums of the squared disparities were explained by the distances in the MDS solutions of the different VCV-conditions. Thus, the two-dimensional perceptual maps reflected the gerbils’ perceptions of the consonants very well.

Similar to the CVCs, the similarity of the perceptual maps of the different VCV-conditions allowed conducting a shared analysis for all VCV-conditions in order to create a generalized perceptual map of the twelve tested consonants in gerbils. Similar clusters as described above for the different VCV-conditions were found for the shared perceptual map of all VCV-conditions and can be seen in Figure 5, panel A. The *DAF* value for the shared perceptual map of all VCV-conditions amounted 89.6%.

**Figure 5:**
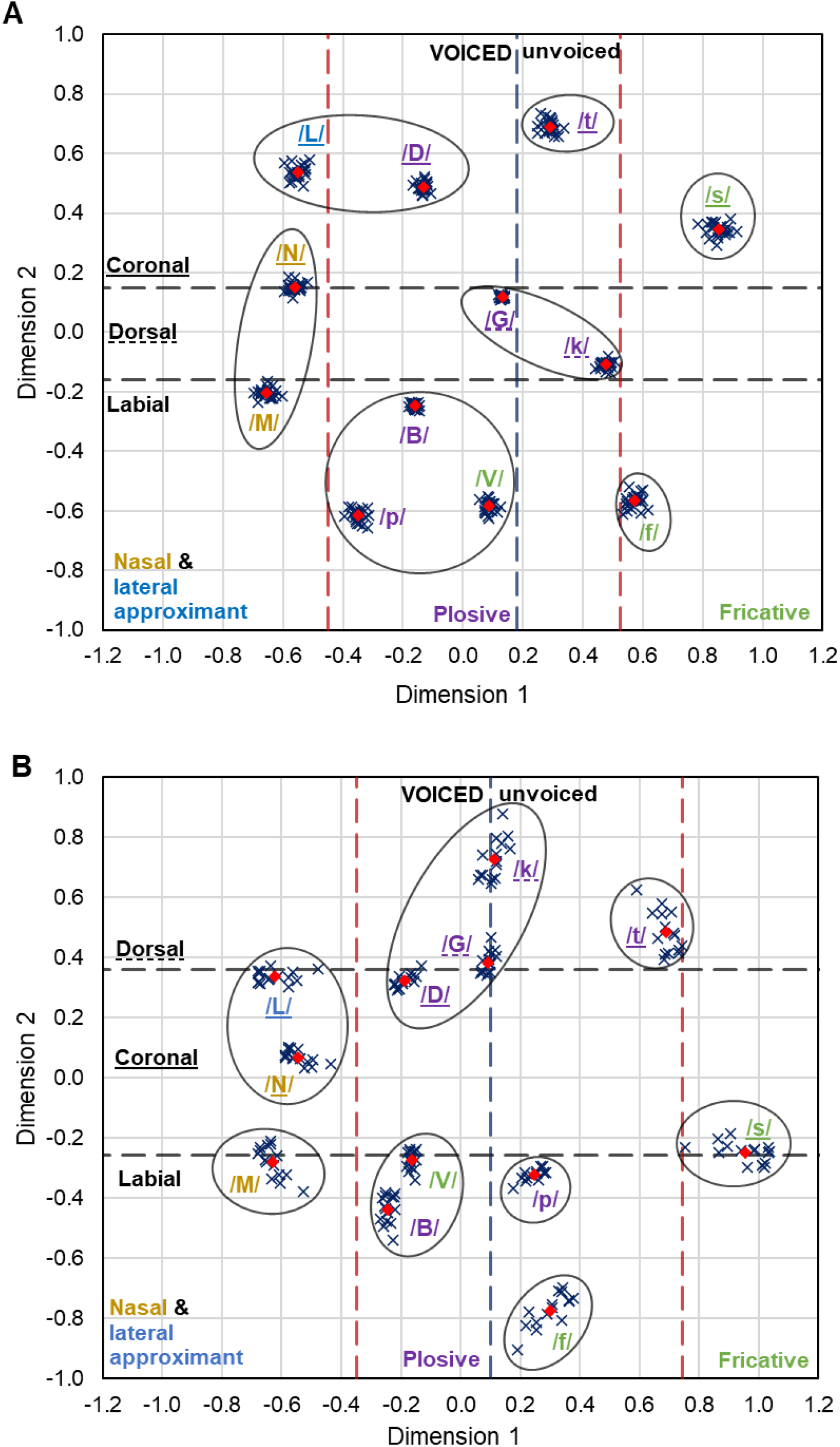
(A): Shared perceptual map of all VCV-conditions for four gerbils. Blue crosses: Individual coordinates for the different VCV-conditions. Red diamonds: Mean coordinates for all VCV-conditions. Black circles: Consonant clusters according to the hierarchical cluster analysis. Dotted lines: Boundaries between the different characteristics of the articulatory features voicing (blue), manner of articulation (red) and place of articulation (black). The different voicing characteristics can be differentiated by case (upper case = voiced, lower case = unvoiced). The place of articulation is marked by underlining (no underline = labial, dotted underline = dorsal, solid underline = coronal). The manner of articulation is indicated by color (purple = plosive, green = fricative, yellow = nasal, blue = lateral approximant). (B): Shared perceptual map of VCVs with the outer vowels /a/, /I/ and /℧/ at -7 dB SNR for five human subjects. Symbols and lines as in (A).

It is evident that the consonants are spatially arranged in a way that the different characteristics of the articulatory features are separated very well from each other. The articulatory feature voicing is split into two groups with the unvoiced consonants on the right side of the perceptual map (except for /p/) and the voiced consonants on the left side. Similarly, for the manner of articulation, the fricative consonants can be found on the right side of the perceptual map (except for /v/), while the nasal consonants and the lateral approximant are situated on the left side. The plosive consonants are arranged in the middle between the other two groups. The different features of the place of articulation are arranged in areas orthogonal to those of the voicing and manner of articulation. The labial consonants are located in the lower area of the perceptual map and the coronal consonants lie in the upper area. In the middle between the labial and the coronal consonants, the dorsal consonants can be found. Thus, the articulatory features of consonants seem to be internally represented along these dimensions in gerbils.

The clusters that were formed during the hierarchical cluster analysis of the human experiments showed many similarities to the clusters of the experiments with the gerbils (see Figure 5, panel B). Similar to the perceptual maps of the gerbils, the consonants /s/, /f/ and /t/ made up clusters on their own and lay distinct from the other consonants in the shared perceptual map for the VCV-conditions of the humans. Further, typical clusters that were found in the perceptual maps of the gerbil VCV-experiments were also present in the human perceptual map and consisted of the dorsal plosives /g/ and /k/ or the voiced labial consonants /b/ and /v/. The consonants within these groups again shared at least two different features for voicing, manner of articulation or place of articulation. In general, as in the perceptual maps of the gerbils, the consonants were spatially arranged in a way that the different characteristics of the articulatory features were separated from each other. Even if the spatial arrangement was rather different for some individual consonants, the overall separation of the different articulatory features was common to the internal representations of both gerbils and humans. This is further supported by a comparison of the perceptual distances between consonant pairs for gerbils and humans in Figure 6 and Figure 7. Even though there was only a weak overall correlation of the distances between the different consonant pairs in the perceptual maps of gerbils and humans (see Figure 6, *R*_2_ = 0.38), significant differences in perceptual distances can be observed depending on similarities of articulatory features for the specific consonant pairs (see Figure 7).

**Figure 6:**
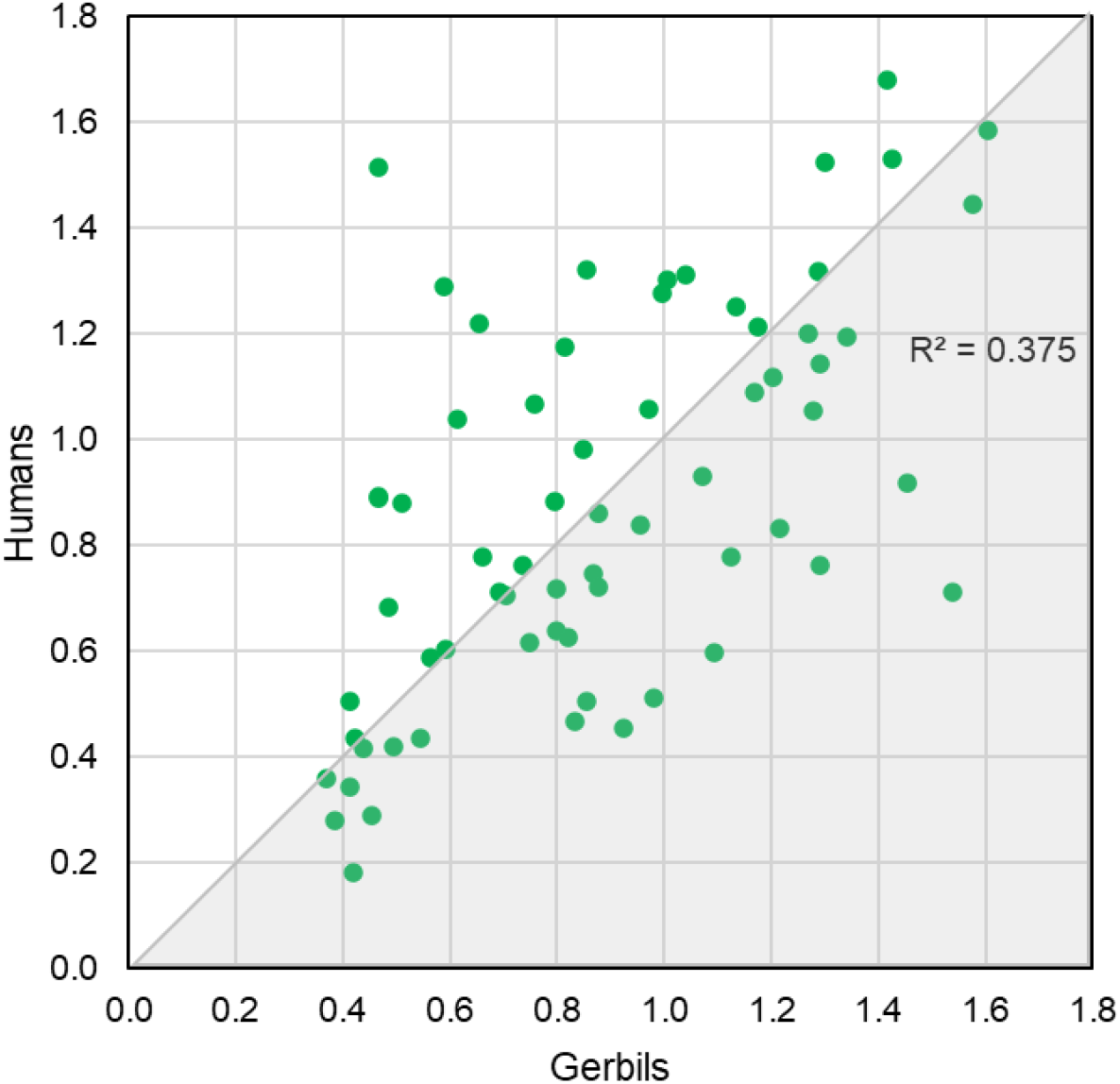
Perceptual distances of consonant pairs for humans vs. gerbils. Perceptual distances from humans plotted against perceptual distances from gerbils for consonant pairs derived from MDS for VCV-conditions.

**Figure 7:**
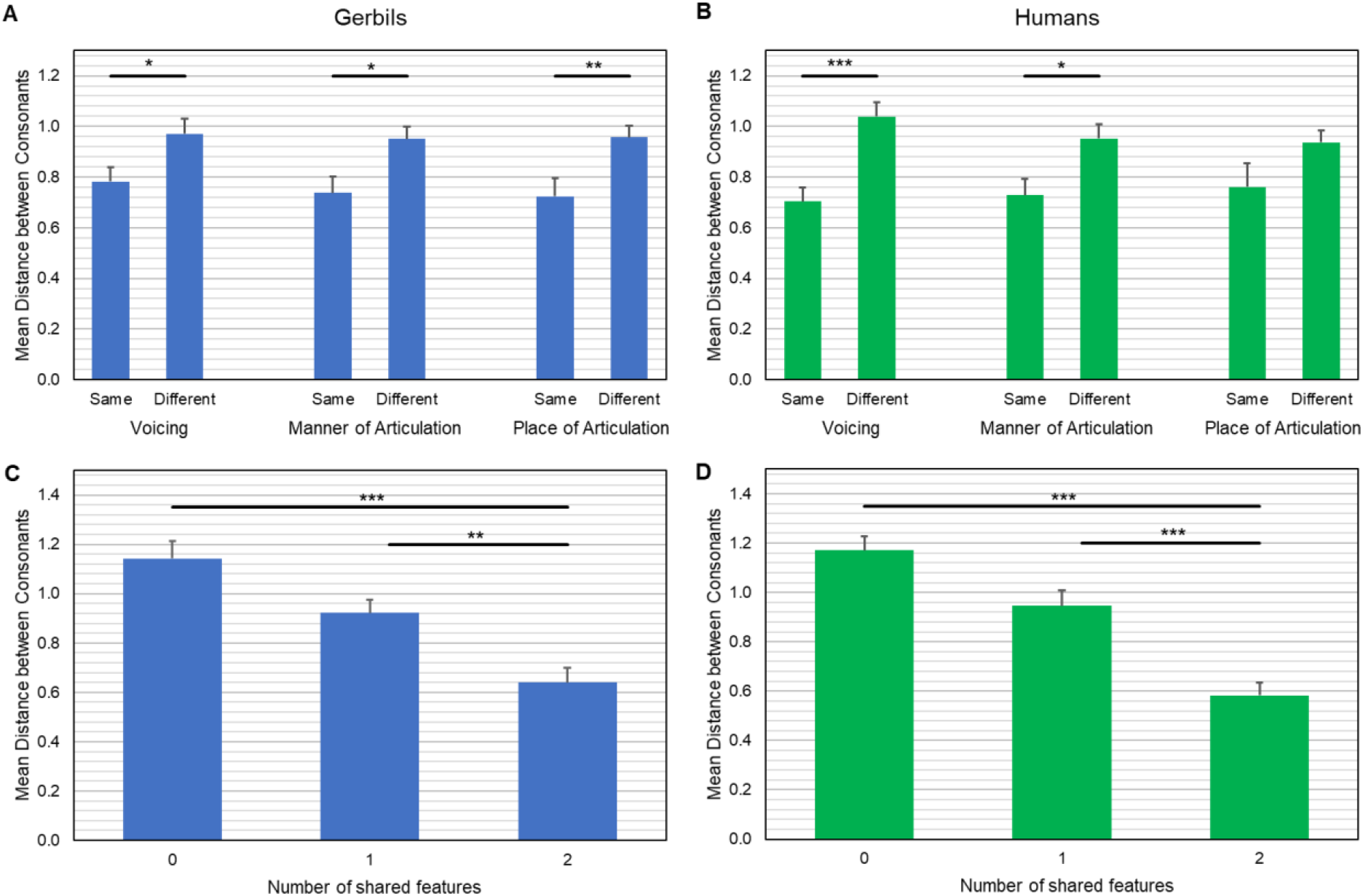
Mean perceptual distances of consonant pairs in relation to differences in voicing, manner, and place of articulation for gerbils (A) and humans (B). Mean perceptual distances of consonant pairs depending on the number of shared articulatory features for gerbils (C) and humans (D). Error bars indicate the standard error of the mean. Significant differences are marked by stars (*: p < 0.05, **: p < 0.01, ***: p < 0.001).

Unpaired two-sample t-tests for the articulatory features voicing (same vs. different), manner of articulation (same vs. different) and place of articulation (same vs. different) showed that differences of the characteristics in either of the three articulatory features led to significantly larger perceptual distances in comparison to consonant pairs with similar characteristics in gerbils (*p*_*Voicing*_ = 0.025, *p*_*Manner*_ = 0.018, *p*_*Place*_ = 0.009; see Figure 7, panel A). In humans, the mean perceptual distance was only significantly larger for consonant pairs that differed in the articulatory features voicing or manner of articulation in comparison to consonant pairs with similar characteristics, but differences in the place of articulation did not lead to significantly larger perceptual distances (*p*_*Voicing*_ < 0.001, *p*_*Manner*_ = 0.020, *p*_*Place*_ = 0.068; see Figure 7, panel B). Moreover, one-way ANOVAs with the number of shared articulatory features (0, 1, 2) as fixed factor and the perceptual distance of the consonant pairs as dependent variable revealed that the more articulatory differences a consonant pair showed, the larger was the perceptual distance between the consonants for gerbils (*F*(2,65) = 13.533, *p* < 0.001; see Figure 7, panel C) and humans (*F*(2,65) = 19.509, *p* < 0.001; see Figure 7, panel D). Post-hoc pairwise comparisons revealed significantly shorter perceptual distances for higher numbers of shared articulatory features within a consonant pair for all comparisons except for an increase from zero to one shared articulatory feature in humans (*p*_0–1_ = 0.056, *p*_0–2_ < 0.001, *p*_1–2_ < 0.001; see Figure 7, panel D) and gerbils (*p*_0–1_ = 0.061, *p*_0–2_ < 0.001, *p*_1–2_ = 0.003; see Figure 7, panel C). This indicates that combinations of the articulatory features are important for the discrimination of consonants.

## 4. Discussion

With the help of the present study, we wanted to investigate how gerbils assess the characteristic features of different speech sounds and if the internal speech representation differs between various logatomes and SNRs. In order to do so, Mongolian gerbils were trained to discriminate between different CVCs and VCVs and perceptual maps of their internal representations of the speech sounds were generated. Finally, the discrimination of vowels and consonants in gerbils was compared to speech sound discrimination in humans.

### 4.1. General Speech Sound Discrimination Abilities in Gerbils and Humans

At large, we saw that the discrimination of vowels is easier for gerbils than the discrimination of consonants (significantly higher *d’* values for CVCs in comparison to VCVs). In line with this, Sinnott and Mosqueda (2003) measured difference limens (DLs) for frequency changes along three synthetic speech continua (vowel, liquid- and stop-consonants) in Mongolian gerbils and found that DLs were lower for the vowel continuum than for the consonant continua. Similarly, humans show higher recognition scores for vowels than for consonants (Jürgens and Brand, 2009; Meyer et al., 2010). Furthermore, vowel recognition is quite robust in humans, while recognition accuracies vary between different consonants (Meyer et al., 2010). Sinnott and Mosteller (2001) also observed - in both gerbils and humans - a facilitated discrimination of vowels compared to the discrimination of consonants.

Despite the similarities between gerbils and humans in their relative ability to discriminate vowels and consonants, the SNR necessary for discriminating both types of speech sounds in humans is markedly below that of gerbils. Even for the ‘easiest’ conditions (CVCs at +15 dB SNR) in the present study, the mean hit rate did not exceed 60% of detected logatome changes. In humans, however, normal-hearing listeners can detect speech sounds with a negligible amount of errors down to -10 dB SPL (Phatak et al., 2008). If the noise level is further increased, the error rate goes from zero to chance performance over a small SNR range (Singh and Allen, 2012). In contrast, the sensitivity index *d’* for gerbils did not differ significantly for an SNR change of 10 dB (+5 dB SNR to +15 dB SNR). Thus, the effect of SNR on the discrimination of speech sounds appears to be different for gerbils and humans. This finding also agrees with previous reports on discrimination of vowels and consonants for gerbils (Eipert and Klump, 2020b; Sinnott et al., 1997; Sinnott and Mosteller, 2001) and humans (Eipert et al., 2019). The saliency for the discrimination of vowels and consonants in gerbils is qualitatively similar to that of humans as indicated by the perceptual maps, but gerbils need higher SNRs for successful speech sound discrimination (Sinnott and Mosteller, 2001).

The reduced overall discrimination ability of both types of speech sounds in gerbils in comparison to humans can be explained by differences in their frequency selectivity. The critical ratio, which is an indirect measure for the frequency selectivity, is higher in gerbils than in humans (Glasberg and Moore, 1990; Kittel et al., 2002; Moore, 2007). In connection to the higher critical ratios, gerbils have larger auditory filter bandwidths than humans, which might correspond to their cochleae being shorter (gerbil cochlea: 11.1 mm (Müller, 1996), human cochlea: 35 mm (Bredberg, 1968)), covering a larger frequency range and having fewer hair cells per critical band (Kittel et al., 2002; Klinge et al., 2010). Thus, formant frequencies can be discriminated better by humans than by gerbils. Apart from a better frequency selectivity or other differences in the general psychoacoustic capacities, human listeners have the large advantage in that they have much more experience with different vowels and consonants, since they have encountered speech tokens of many different talkers at different SNRs and in different reverberations throughout their lifetime (Jürgens et al., 2014), whereas the gerbils were trained with human speech sounds only for several months for the purpose of this study. Moreover, speech is an essential component in many aspects of everyday human life, increasing the importance of the ability to successfully discriminate speech sounds for humans.

Despite their quantitatively lower sensitivity for discriminating human speech sounds, gerbils demonstrated an ability to discriminate between different vowels and consonants, which further implies that the phonetic boundary of these speech sounds is a natural, physiological one that is also exhibited by nonhuman animals (Kuhl and Miller, 1975). Thus, gerbils are a promising small-mammal model for the investigation of higher-level processes of speech perception (Sinnott and Mosteller, 2001).

### 4.2. Discrimination of Vowels

Beside the general comparison of speech sound discrimination in gerbils and humans, the perceptual maps were analyzed with respect to the stimulus features leading to salient differences between the different logatomes in gerbils. Vowels appear to be internally represented according to the frequencies of their first and second formant, which changed along the two dimensions of the perceptual maps. In addition to the spectral features of vowels, also their articulatory features corresponded to the two dimensions of the perceptual maps. The frequencies of F1 and F2 reflect the tongue height and the position of the tongue articulation (Sinnott et al., 1997), respectively. Thus, not only spectral but also articulatory features of the vowels could be assigned to the dimensions of the perceptual maps. This is also in line with findings from Mesgarani et al. (2008), who observed that those vowels with back places of articulation (/u:/, /o:/, /ɔ/, /℧/) show rather concentrated low frequency activities, and this single frequency peak broadens and splits up for more central vowels (/a/, /a:/) until two separated peaks spaced over a larger range of frequencies can be observed for front vowels (/ε/, /I/, /e:/, /i:/). These peaks represent the spectral pattern of the formants in the different vowels (Ladefoged and Johnson, 2011), corresponding to the perceptual clusters in the present study. Regarding the relative importance of the discrimination cues, the results shown in Figure 4 demonstrate that the tongue position (corresponding to F2) is of higher relevance for vowel discrimination than the tongue height (corresponding to F1), since differences in tongue height did not lead to significantly larger perceptual distances in contrast to differences in tongue position.

This close relation between the perceptual maps and both spectral and articulatory features, and the cues underlying the discrimination of vowels are common to gerbils and humans. Thus, irrespective of the different SNRs that were used for humans and gerbils and the lower salience in discriminating human speech sounds for gerbils, the perception of different vowels appears to be qualitatively similar for humans and gerbils, suggesting that the mechanisms underlying this discrimination might be the same in gerbils and humans.

In 2009, Jürgens and Brand used the same stimuli in an experimental procedure in which the different logatomes had to be recognized and indicated on a list by human subjects. Consistent with the vowel clusters in the present study, they found that the CVCs with the central vowels /ɔ/, /o:/, /℧/ and /u:/ were those with the largest numbers of confusions. The especially high confusion of the central vowels /o:/ and /u:/ may be due to their average spectra being most effectively masked by the speech-shaped noise (Meyer et al., 2007). Furthermore, Jürgens and Brand (2009) observed a substantial confusion between the vowels /a/ and /a:/ which fits well with their very short perceptual distances in the perceptual maps of the present study. In line with this, also Meyer et al. (2010) found the highest error rates in discriminating the vowel pair /a/ and /a:/ and explained this with their phonetic and spectral similarity. In contrast to the other vowels in the stimulus set that differ in tongue height and position, /a/ and /a:/ only vary with respect to their duration (Meyer et al., 2010).

Besides the discriminability of vowels, also the identification of important features for discrimination was addressed in previous studies, however, with varying outcomes. As in the present study, the formant frequencies or quantities derived from different formants were already considered as suitable equivalences for the different perceptual dimensions. Ohl and Scheich (1997) investigated the cortical representation of formant frequencies focusing on the first two formants. They suggested that the frequency distance between F1 and F2 might be a potentially useful parameter for vowel identification since this difference has a neuronal representation in A1. This distance between F1 and F2 may be more robust against speaker variations than F1 or F2 alone. However, in the present study in both gerbils and humans the pure frequencies of F1 and F2 were correlated more strongly with either the first or the second dimension of the perceptual maps than the frequency difference between these two formants.

In a two-dimensional perceptual map of vowel clusters shown in the study from Phatak and Allen (2007), the dimensions were found to be closely related to the vowel duration and F2 frequency, respectively. Blamey et al. (1987) categorized the vowels by duration assigning /a(:)/, /i:/, /ɔ/ and /u:/ to the long-duration category and /ε/, /I/ and /℧/ to the short-duration category. This categorization of vowels according to their duration is not represented along any of the two dimensions of the perceptual maps in the present study. Thus, the finding that one dimension of a two-dimensional projection of vowels is related to the vowel duration (Phatak and Allen, 2007) cannot be confirmed for the present sample of logatomes. The study from Blamey et al. (1987) also categorized the different vowels according to their F1 and F2 frequencies. These categories for F1 (high: /a(:)/, mid: /ε/, /ɔ/, /℧/, low: /I/, /i:/, /u:/) and F2 (high: /ε/, /I/, /i:/, mid: /a(:)/, /u:/, low: /ɔ/, /℧/) (Blamey et al., 1987) are reflected much better than the vowel duration along the second and first dimension in the perceptual maps of the present study, respectively.

The preceding and subsequent consonants in the logatomes only had a minor influence on the discrimination of the central vowels corresponding to a previous report of confusion patterns of CV-stimuli in humans (Phatak and Allen, 2007). Further, previous vowel studies with different animal species found that these animals (e.g. chinchillas and budgerigars) were able to generalize within vowel categories in spite of variations introduced by different talkers (Burdick and Miller, 1975; Dooling and Brown, 1990). This can be confirmed for gerbils as well, since they grouped the same vowels spoken by different speakers into one category.

In summary, the frequencies of F1 and F2 being determined by the manner and place of articulation are most important for the discrimination of vowels in gerbils. Most importantly, the perception of the different human vowels appears to be very similar in gerbils and humans.

### 4.3. Discrimination of Consonants

The interpretation of the perceptual maps for the VCV-conditions was less straightforward than for the CVC-conditions, since consonants in contrast to vowels are not characterized by clear formants that provide spectral features for their discrimination. Thus, the dimensions in the perceptual maps of the VCV-conditions were interpreted solely based on the articulatory features of the consonants. The dashed lines in Figure 5 indicated approximate boundaries between the different characteristics of the articulatory features separating these well from each other. Thus, in gerbils the articulatory features voicing, manner and place of articulation seem to be perceptually represented along these two dimensions leading to short perceptual distances for consonants with the same articulatory characteristics.

Also in the perceptual map of the human subjects, the logatomes were arranged in a way that consonants with the same articulatory characteristics showed spatial proximity. The overall separation of the different characteristics of the articulatory features was preserved and common to the perceptual maps of both gerbils and humans, although the location of individual consonants could differ. Consequently, differences in combinations of articulatory features led to large perceptual distances and low confusion probabilities. This supports the hypothesis, that differences in combinations of the articulatory features are important for the discrimination of consonants for both gerbils and humans. One difference between gerbils and humans, however, might be the relative importance of the different articulatory features for consonant discrimination. Figure 7 shows that different places of articulation did not lead to significantly larger perceptual distances in humans, whereas differences in this articulatory feature were the most important cue for consonant discrimination in gerbils. Differences in voicing were the most important cue for consonant discrimination in humans and less important in gerbils. Moreover, in humans already one articulatory difference led to significantly larger perceptual distances in comparison to zero articulatory differences while two differences were needed in gerbils. This might be a consequence of the lower sensitivity for discriminating logatomes by gerbils compared to humans.

Miller and Nicely (1955) observed in humans that for low SNRs consonants form three basic clusters of confusable sounds, namely unvoiced consonants, voiced (non-nasal) consonants and nasal consonants. For higher SNRs, they found that both the unvoiced and the voiced (non-nasal) group of consonants split up into subgroups of plosive and fricative consonants. Thus, for higher SNRs, they basically made the same observations as we did in the present study for gerbils and humans. Although Miller and Nicely (1955) did not consider the place of articulation, their results are in accordance with voicing and manner of articulation playing a pivotal role in the discrimination of consonants. Resembling the results of the present study, Mesgarani et al. (2008) found perceptual and neural consonant categories in ferrets that were defined according to the manner of articulation. Ferrets distinguished between the plosives /p/, /t/ and /k/, the fricatives /f/ and /s/ and the nasals /m/ and /n/. Consonants of the same category were found to be more confusable than consonants of different categories.

Another study that made use of the same stimulus set and masker as in the present experiments found that the highest recognition scores for consonants in humans can be observed for /s/ and /t/ (Meyer et al., 2010). It was suggested that the differences in recognition rates can be explained mainly by differences in the long-term spectra of speech and noise (Jürgens and Brand, 2009), since the spectra for /s/ and /t/ differ more from the masking-noise spectrum than the other consonants. This can also explain that /s/ and /t/ stood out of the other consonants in the perceptual maps of the present study by mostly making up clusters on their own and by exhibiting the largest inter-consonant distances.

Additionally, the voice-onset time (VOT) may be important for the discrimination of plosive consonants, being characterized by their sudden and spectrally broad onset (Mesgarani et al., 2008). Voiced (/b/, /d/, /g/) and unvoiced (/p/, /t/, /k/) plosives can be differentiated by a timing difference between the onset of the plosive burst and the onset of voicing within the consonant (Lisker and Abramson, 1964). For humans, the VOT is an important cue in many languages to signal the articulatory feature voicing (Cho and Ladefoged, 1999; Lisker and Abramson, 1964; Toscano and Lansing, 2019). In the present study, not only humans but also gerbils were able to distinguish the voiced from the unvoiced plosives quite well. This suggests that also gerbils are able to make use of differences in VOT as a cue for voicing in order to discriminate between plosive consonants.

In summary, the different articulatory features seem to play a pivotal role in the discrimination of consonants in gerbils as well as in humans. Thus, differences in combinations of the features voicing, manner of articulation and place of articulation strongly influence the confusion probabilities of consonants.

## 5. Conclusion

In conclusion, we found that gerbils can discriminate more easily between vowels than between consonants. Vowels were discriminated predominantly based on differences in spectral features of their formants. These spectral features are determined by the place and manner of articulation, which are then reflected in the frequency of F2 and F1, respectively. The discrimination of consonants mostly depends on the combination of their articulatory features (voicing, manner of articulation and place of articulation). Further, the basic mechanisms of speech sound discrimination seem to be very similar in gerbils and humans. However, the overall sensitivity for discriminating human speech sounds differs between gerbils and humans. Gerbils proved to be a suitable animal model for investigating the discrimination of human speech sounds. Further studies with aged gerbils may reveal the effect of age-related hearing loss on speech sound discrimination and the underlying mechanisms. This may provide for explaining speech comprehension deficits in elderly human listeners.

## Author Contributions

GMK and CJ conceived the idea and designed the study. CJ collected the data. CJ and RB analyzed the data. CJ, GMK, and RB interpreted the data. CJ drafted the manuscript. GMK, CJ, and RB critically revised the manuscript.

## CRediT Authorship Contribution Statement

**Carolin Jüchter:** Conceptualization, Formal analysis, Investigation, Data curation, Writing - Original draft, Visualization.

**Rainer Beutelmann:** Software, Formal analysis, Data curation.

**Georg M. Klump:** Conceptualization, Writing - Review & Editing, Supervision.

## Acknowledgements

We thank Melissa Jäger for her contribution to data collection. Further, we thank Dawid Fandrich, Franziska Berger, Laura-Janine Döring, Melissa Jäger and Nadine Dyszkant for their participation in the study as part of a student course. This work was funded by the Deutsche Forschungsgemeinschaft (DFG, German Research Foundation) under Germany’s Excellence Strategy – EXC 2177/1 - Project ID 390895286).

## Supplementary Material

**Supplementary Material A:**
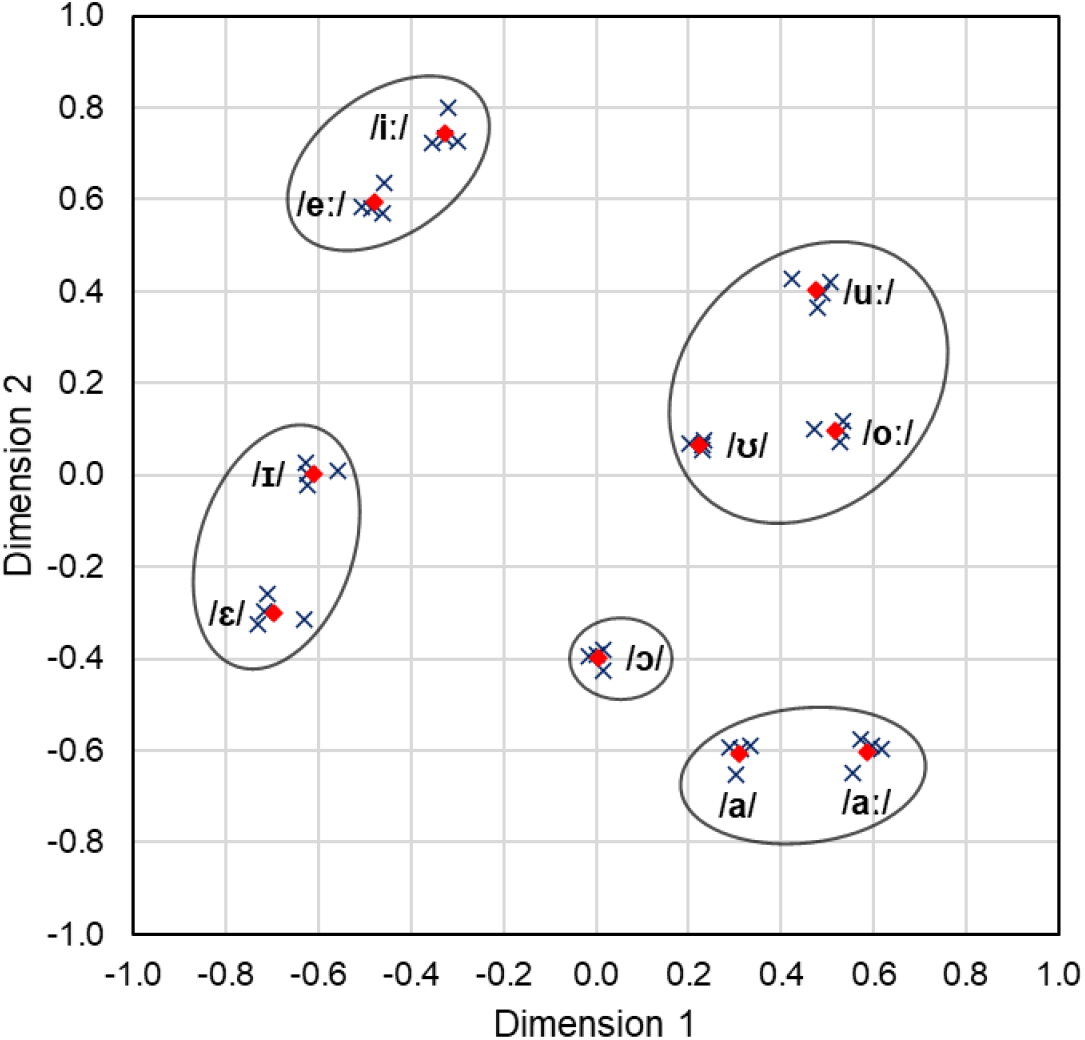
Perceptual map of the CVC-condition with the outer consonant /b/ at +5 dB SNR. Blue crosses: Perceptual vowel coordinates of the different individuals. Red diamonds: Mean perceptual vowel coordinates of all individuals. Black circles: Vowel clusters according to the hierarchical cluster analysis.

**Supplementary Material B:**
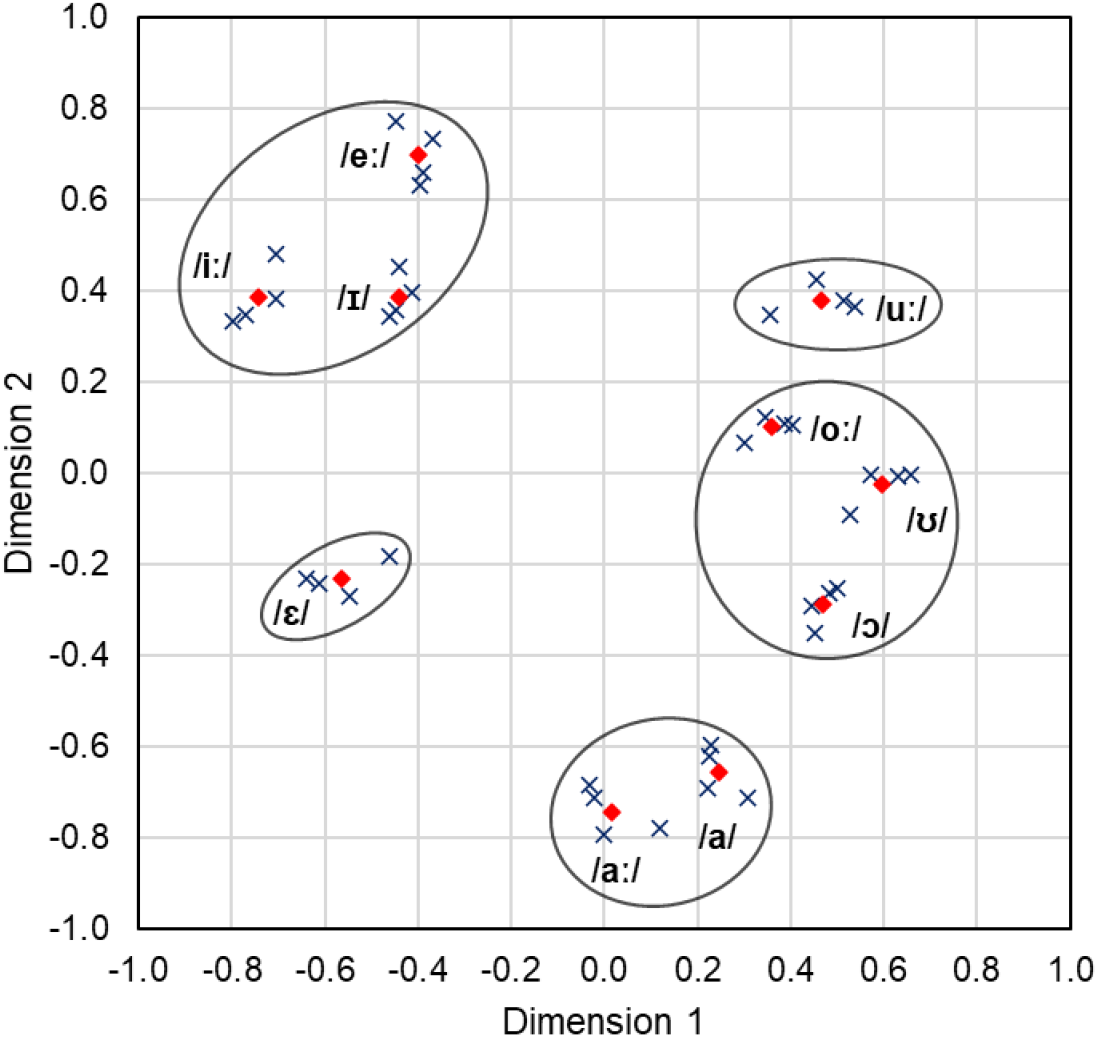
Perceptual map of the CVC-condition with the outer consonant /b/ at +15 dB SNR. Blue crosses: Perceptual vowel coordinates of the different individuals. Red diamonds: Mean perceptual vowel coordinates of all individuals. Black circles: Vowel clusters according to the hierarchical cluster analysis.

**Supplementary Material C:**
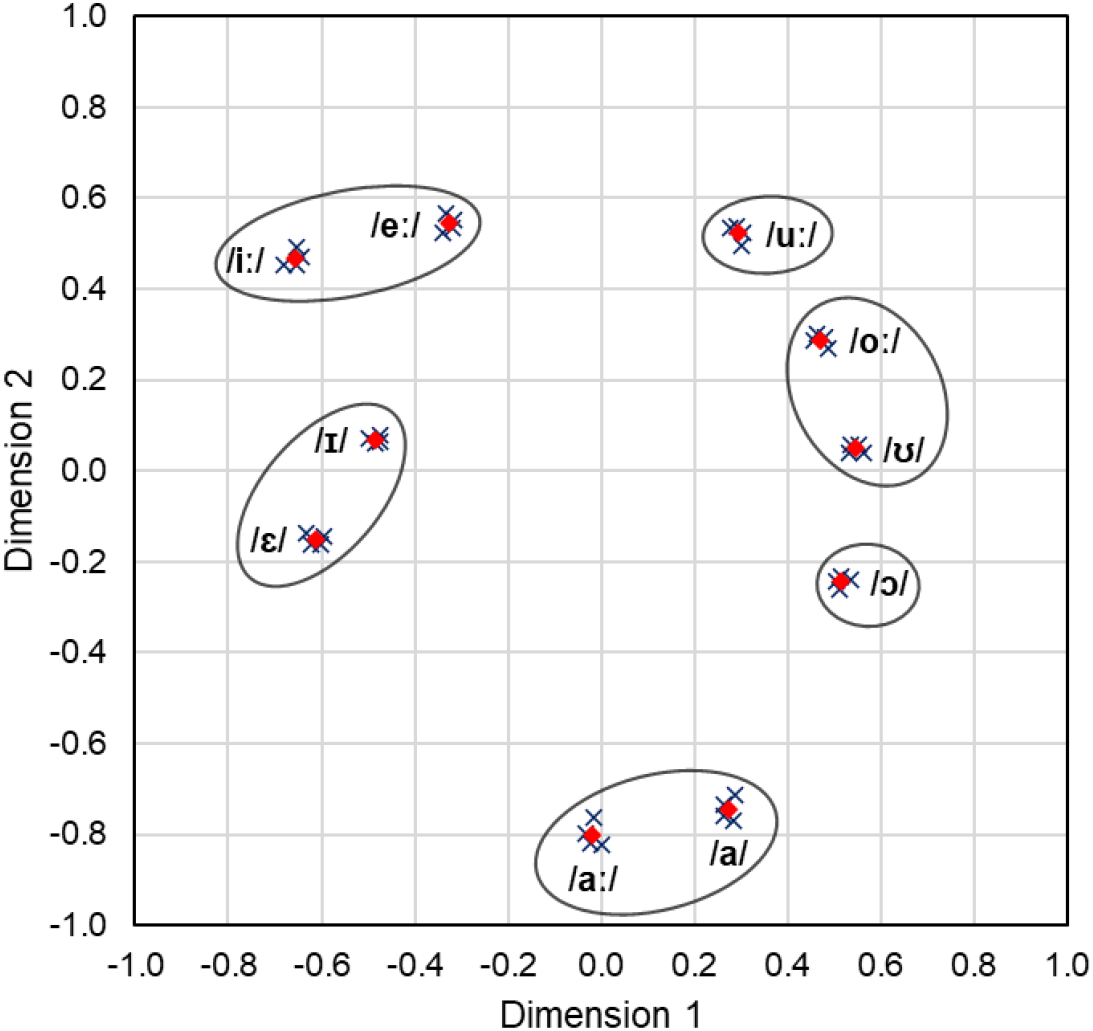
Perceptual map of the CVC-condition with the outer consonant /d/ at +5 dB SNR. Blue crosses: Perceptual vowel coordinates of the different individuals. Red diamonds: Mean perceptual vowel coordinates of all individuals. Black circles: Vowel clusters according to the hierarchical cluster analysis.

**Supplementary Material D:**
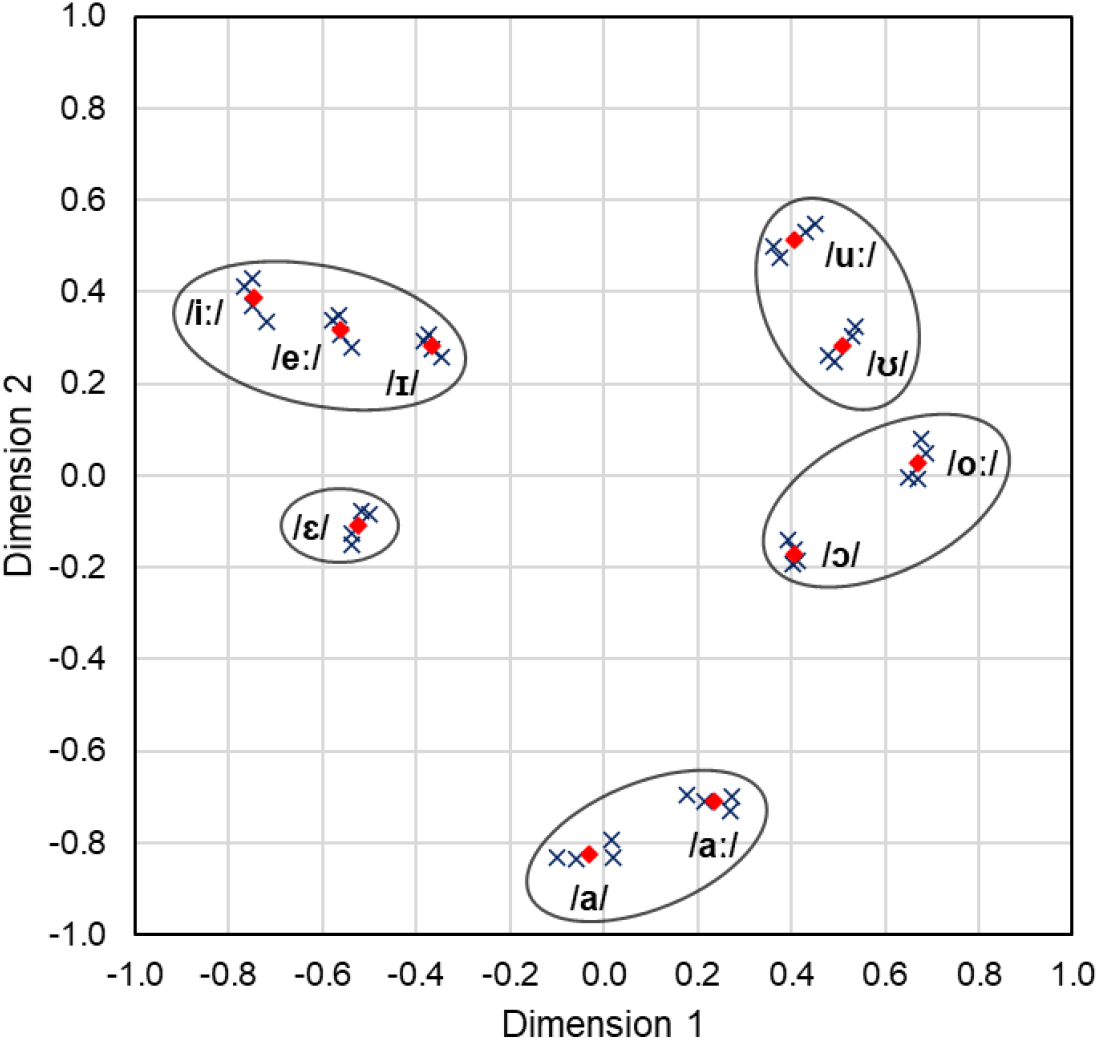
Perceptual map of the CVC-condition with the outer consonant /d/ at +15 dB SNR. Blue crosses: Perceptual vowel coordinates of the different individuals. Red diamonds: Mean perceptual vowel coordinates of all individuals. Black circles: Vowel clusters according to the hierarchical cluster analysis.

**Supplementary Material E:**
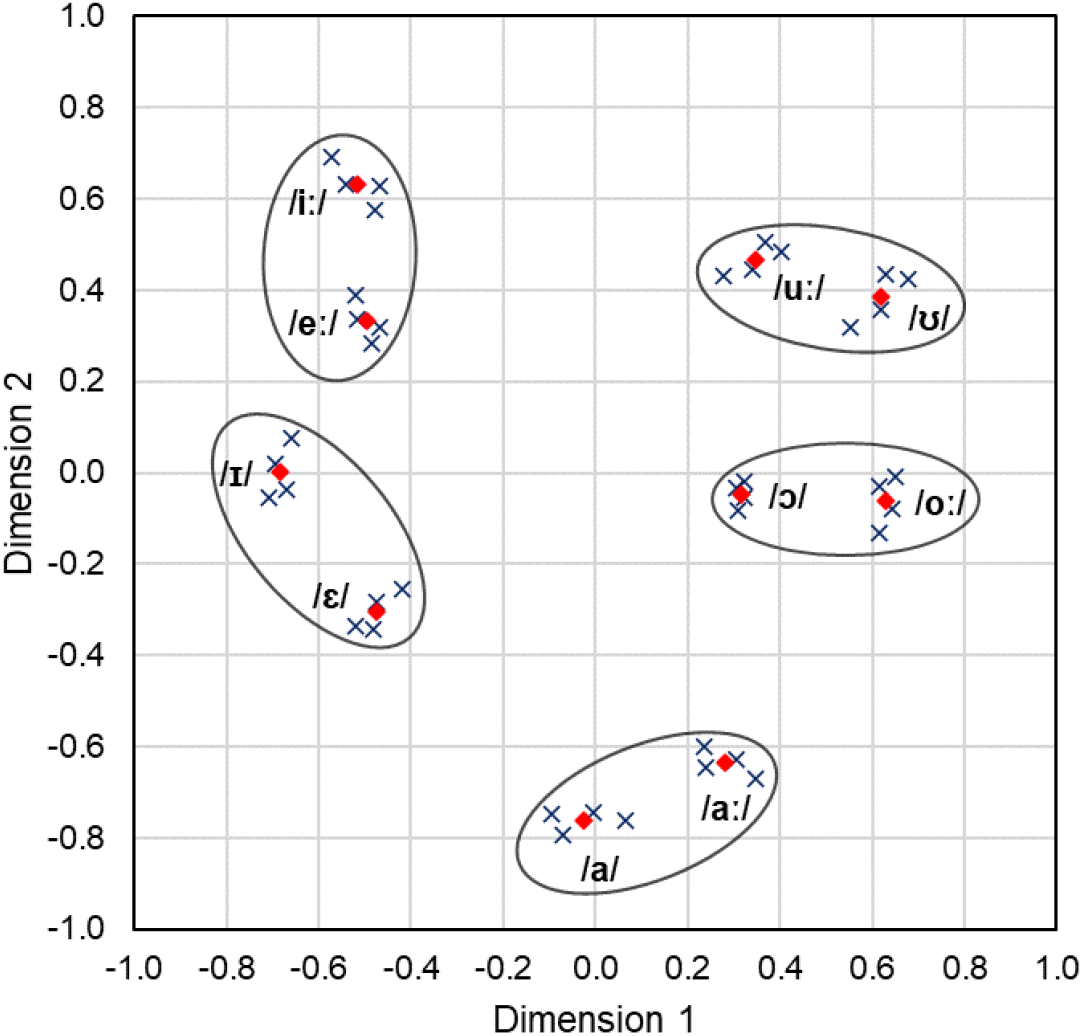
Perceptual map of the CVC-condition with the outer consonant /s/ at +5 dB SNR. Blue crosses: Perceptual vowel coordinates of the different individuals. Red diamonds: Mean perceptual vowel coordinates of all individuals. Black circles: Vowel clusters according to the hierarchical cluster analysis.

**Supplementary Material F:**
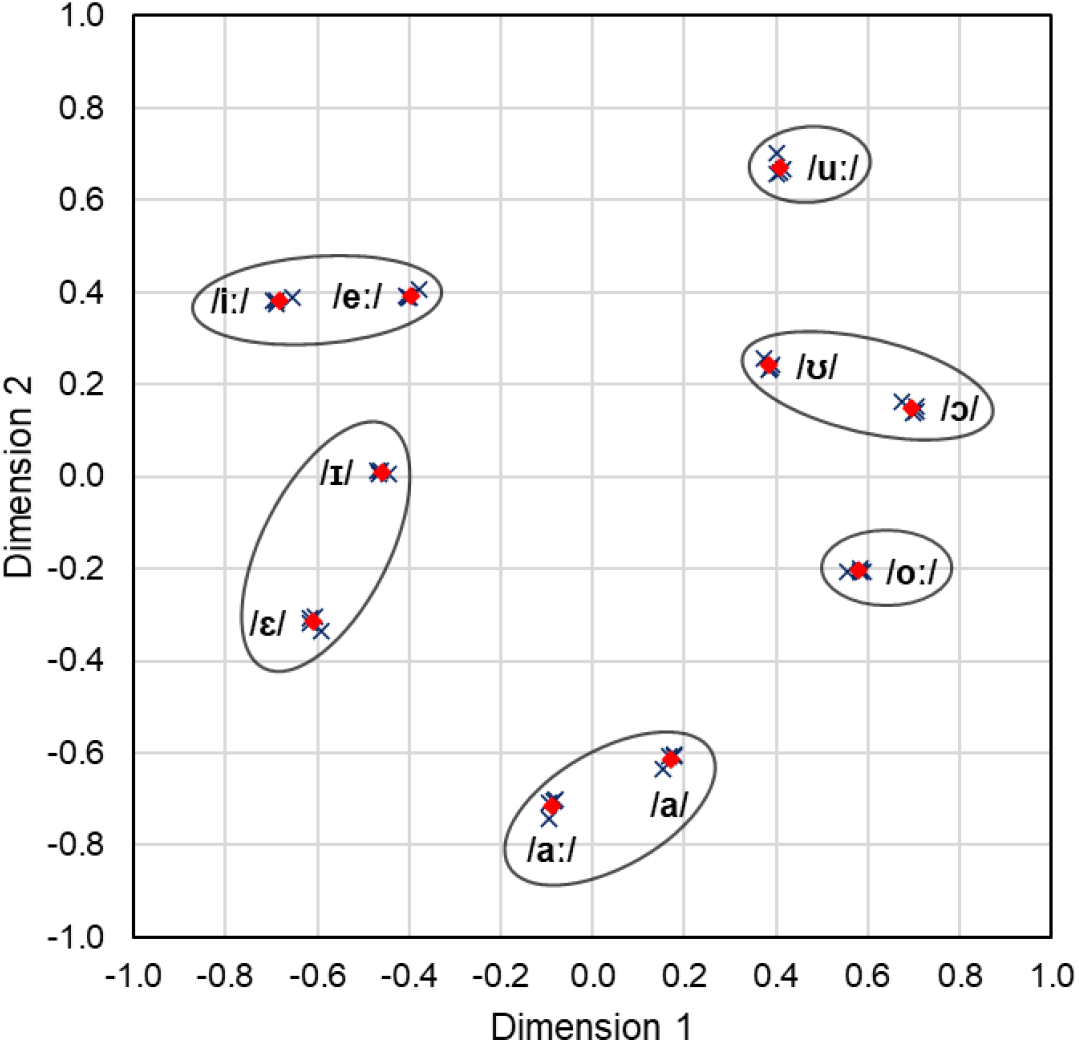
Perceptual map of the CVC-condition with the outer consonant /s/ at +15 dB SNR. Blue crosses: Perceptual vowel coordinates of the different individuals. Red diamonds: Mean perceptual vowel coordinates of all individuals. Black circles: Vowel clusters according to the hierarchical cluster analysis.

**Supplementary Material G:**
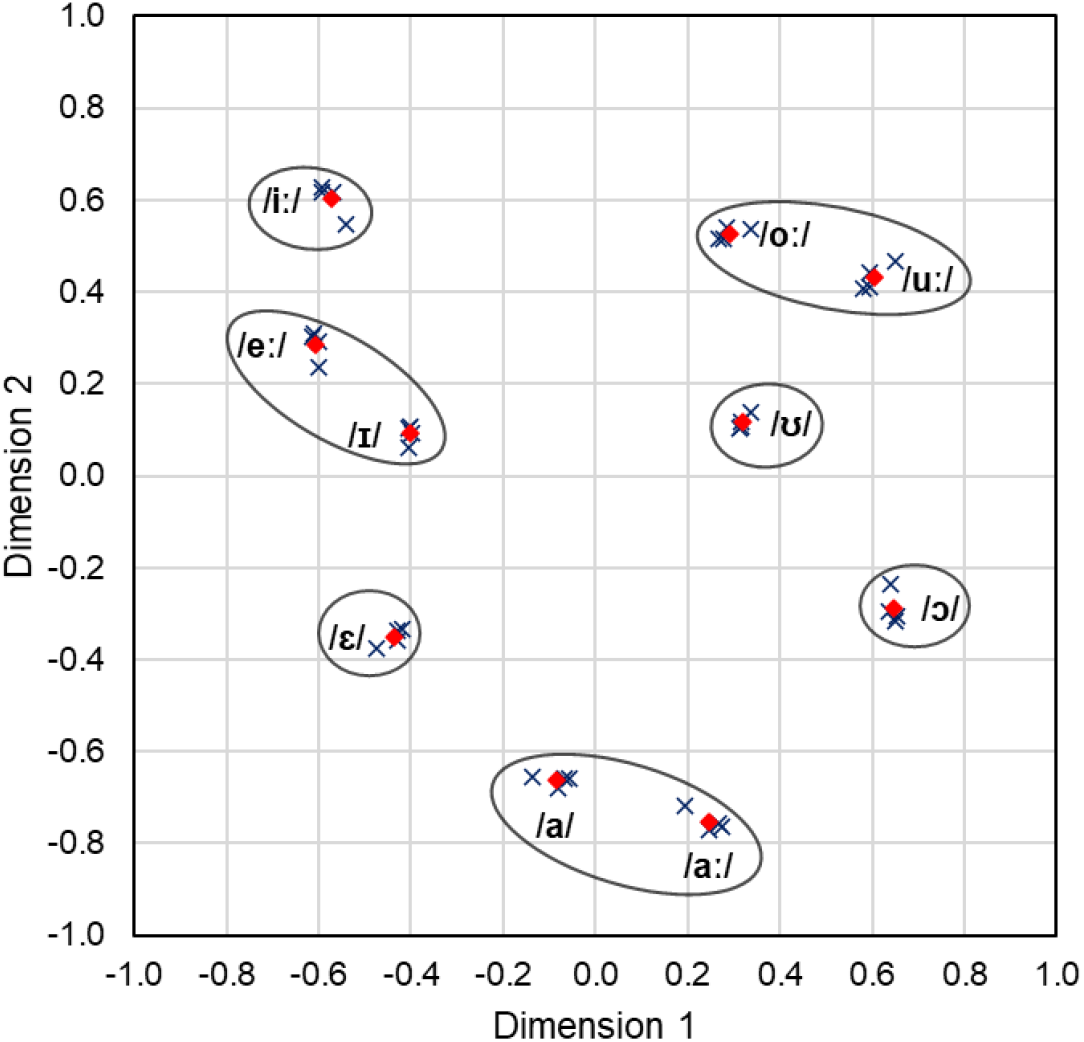
Perceptual map of the CVC-condition with the outer consonant /t/ at +5 dB SNR. Blue crosses: Perceptual vowel coordinates of the different individuals. Red diamonds: Mean perceptual vowel coordinates of all individuals. Black circles: Vowel clusters according to the hierarchical cluster analysis.

**Supplementary Material H:**
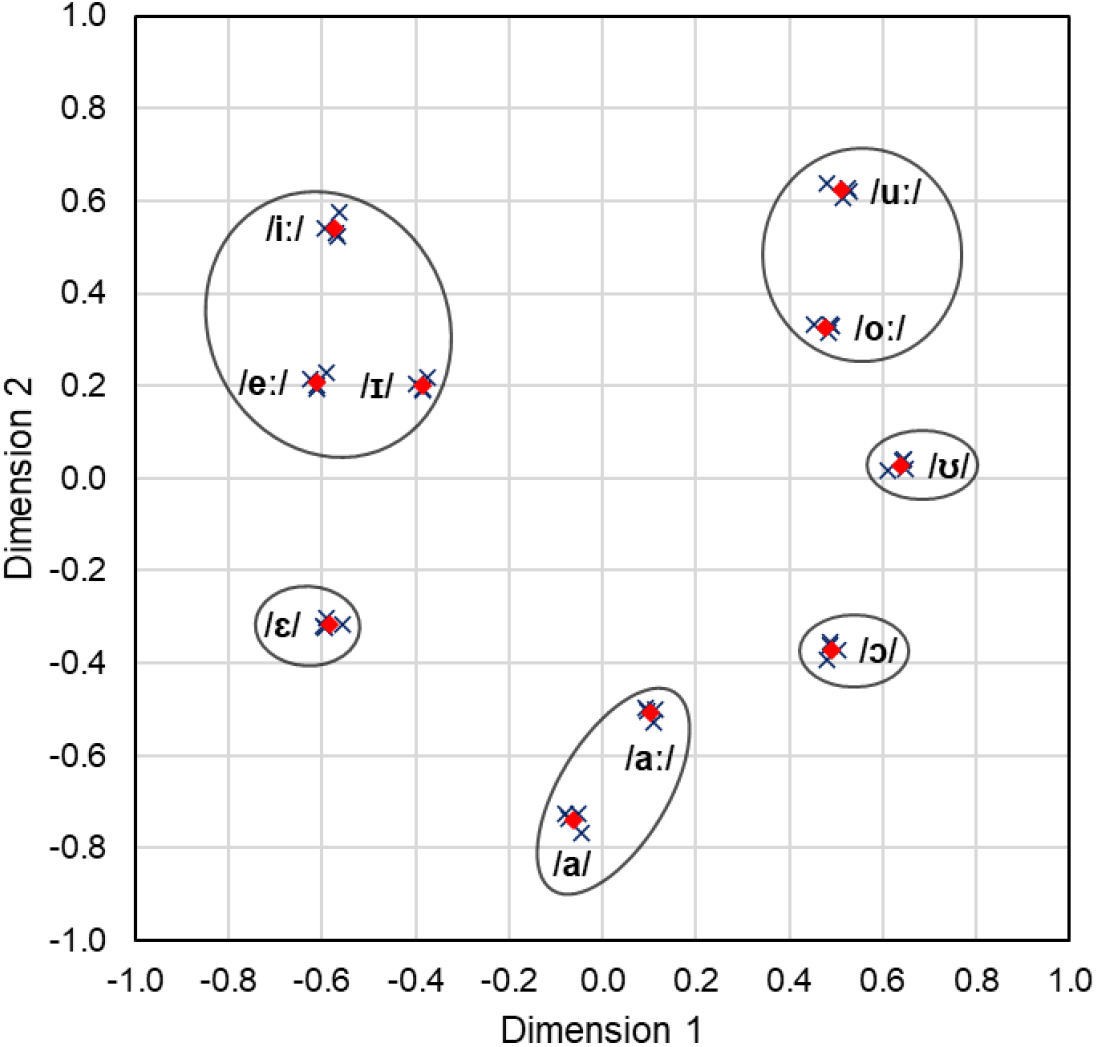
Perceptual map of the CVC-condition with the outer consonant /t/ at +15 dB SNR. Blue crosses: Perceptual vowel coordinates of the different individuals. Red diamonds: Mean perceptual vowel coordinates of all individuals. Black circles: Vowel clusters according to the hierarchical cluster analysis.

**Supplementary Material I:**
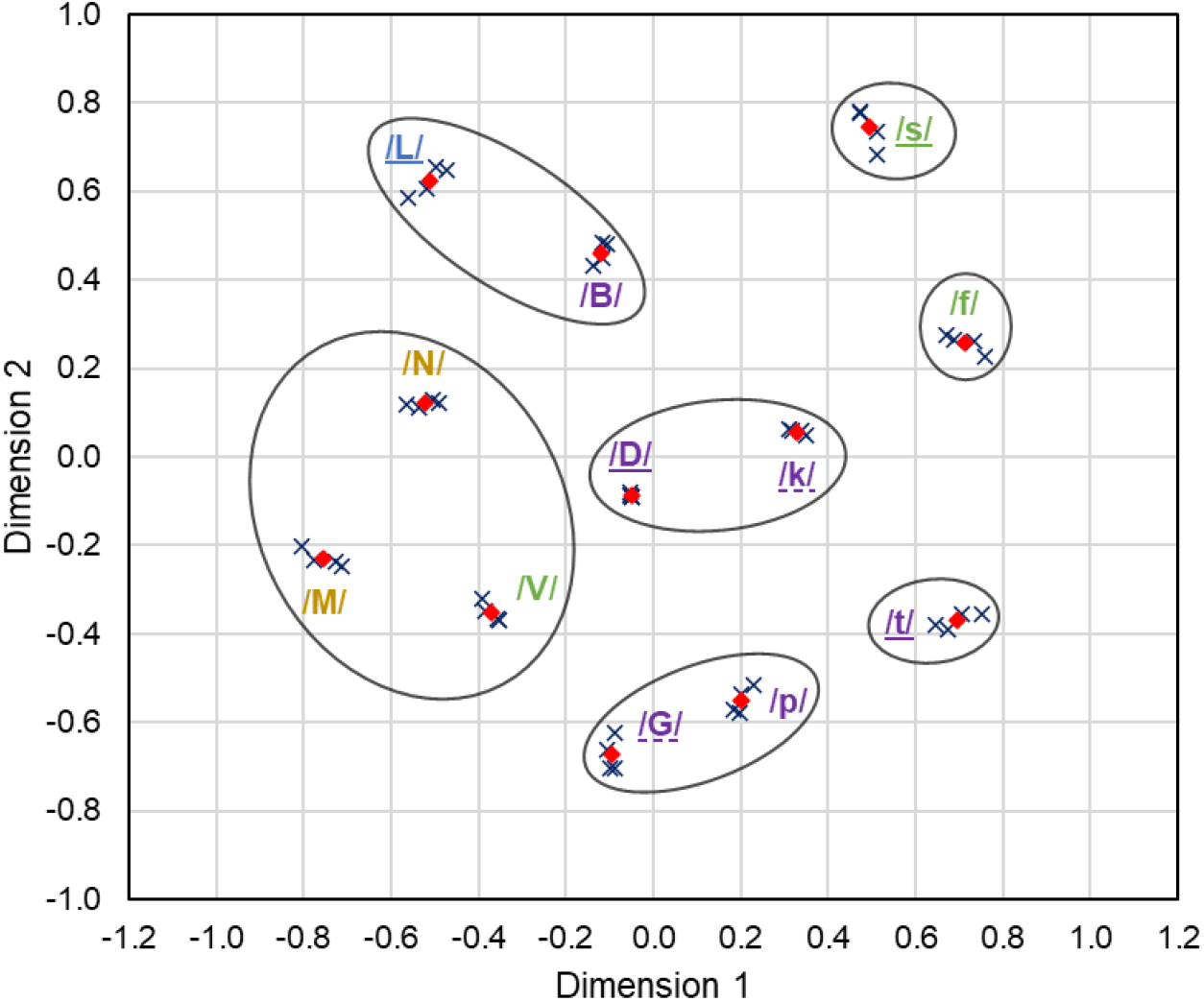
Shared perceptual map of the VCV-condition with the outer vowel /a/ at +5 dB SNR. Blue crosses: Perceptual consonant coordinates of the different individuals. Red diamonds: Mean perceptual consonant coordinates of all individuals. Black circles: Consonant clusters according to the hierarchical cluster analysis.

**Supplementary Material J:**
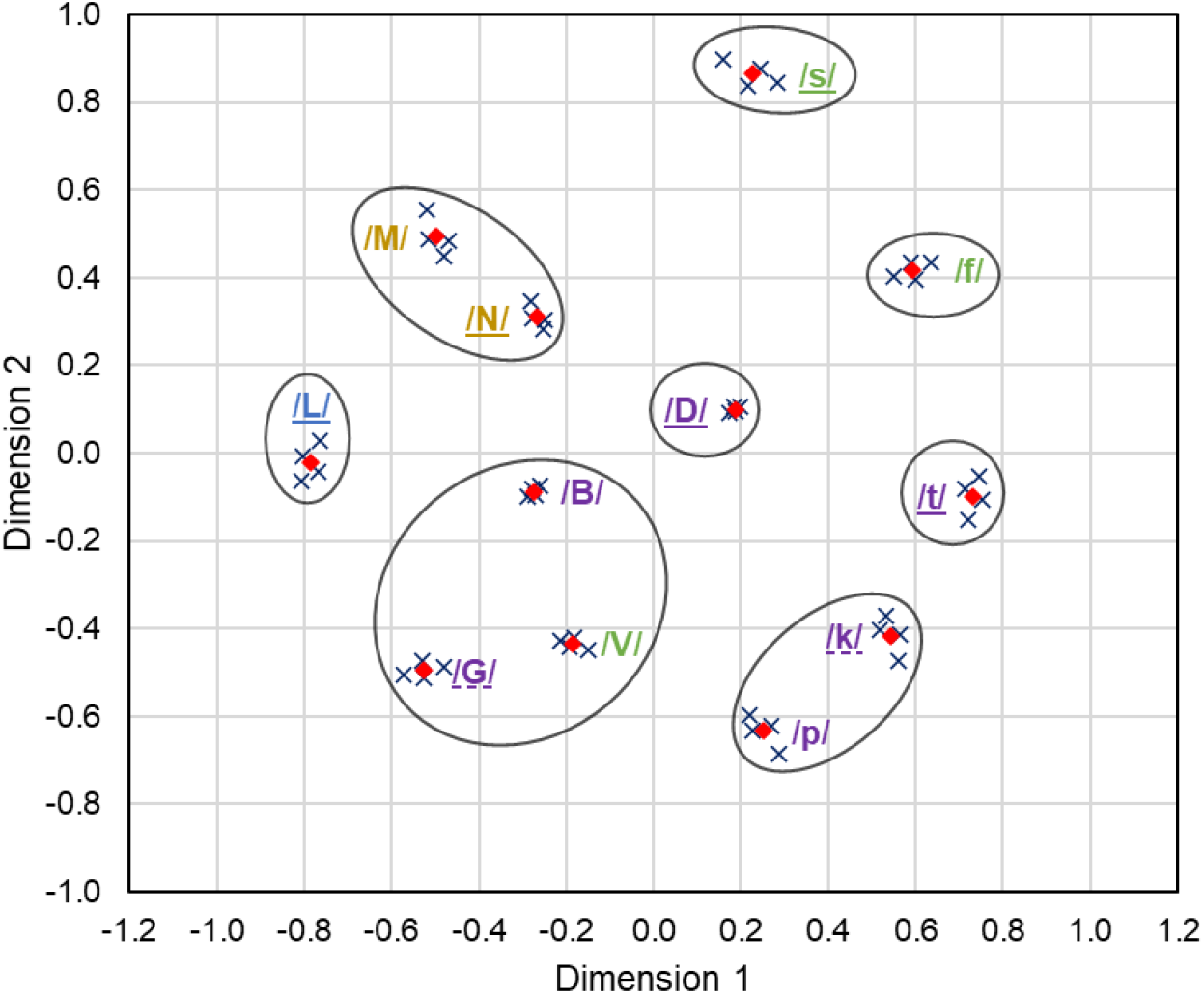
Shared perceptual map of the VCV-condition with the outer vowel /a/ at +15 dB SNR. Blue crosses: Perceptual consonant coordinates of the different individuals. Red diamonds: Mean perceptual consonant coordinates of all individuals. Black circles: Consonant clusters according to the hierarchical cluster analysis.

**Supplementary Material K:**
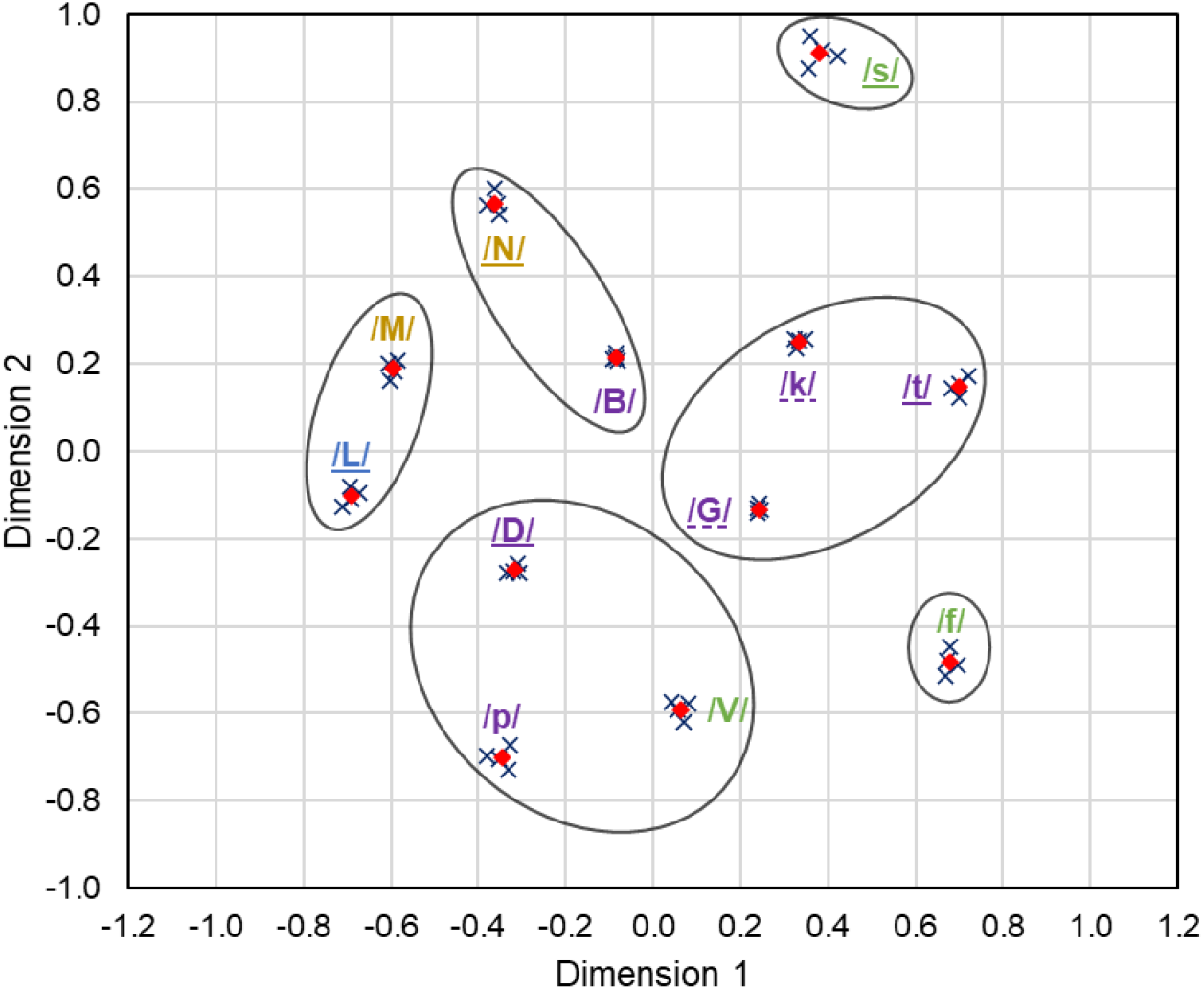
Shared perceptual map of the VCV-condition with the outer vowel /I/ at +5 dB SNR. Blue crosses: Perceptual consonant coordinates of the different individuals. Red diamonds: Mean perceptual consonant coordinates of all individuals. Black circles: Consonant clusters according to the hierarchical cluster analysis.

**Supplementary Material L:**
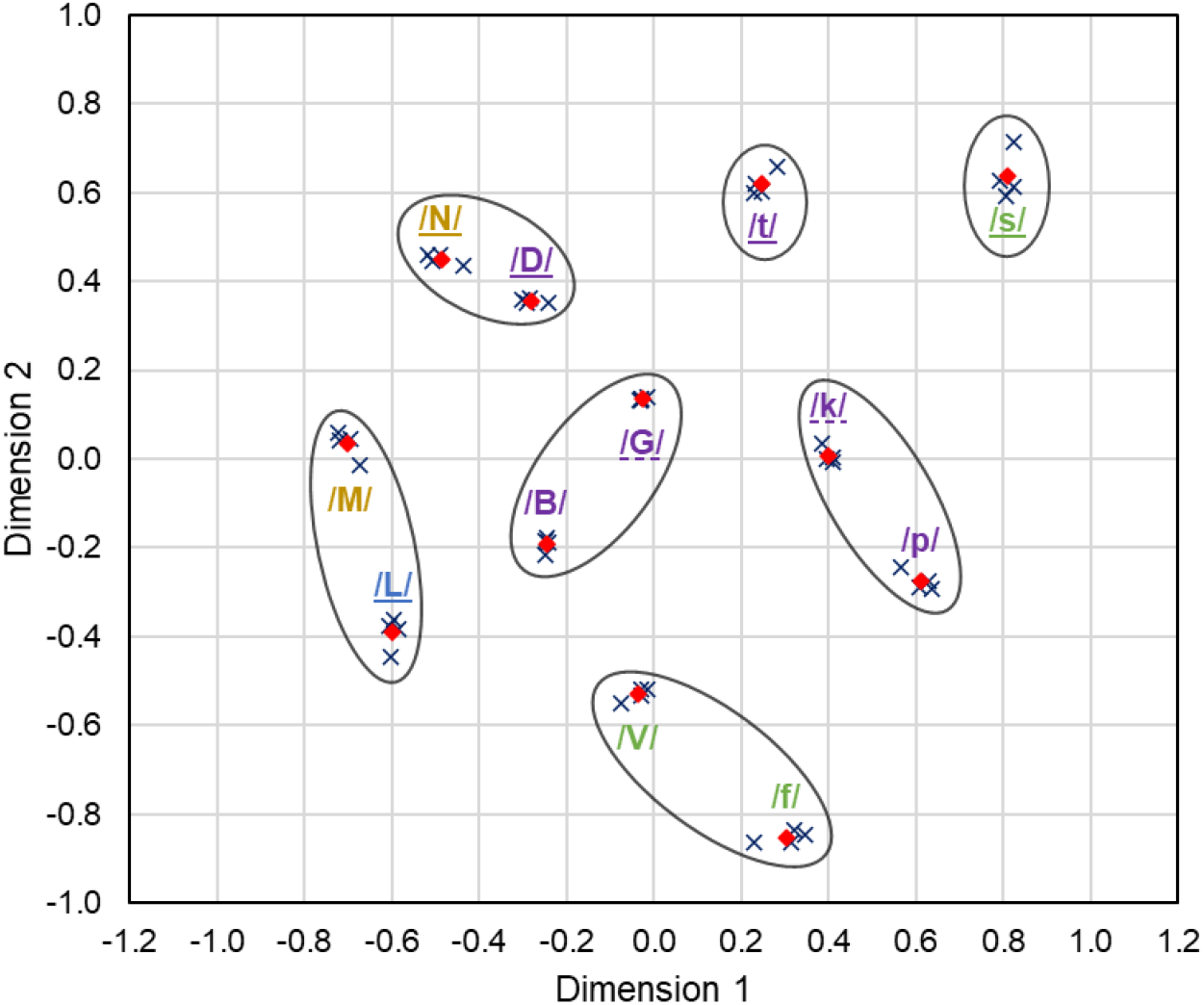
Shared perceptual map of the VCV-condition with the outer vowel /I/ at +15 dB SNR. Blue crosses: Perceptual consonant coordinates of the different individuals. Red diamonds: Mean perceptual consonant coordinates of all individuals. Black circles: Consonant clusters according to the hierarchical cluster analysis.

**Supplementary Material M:**
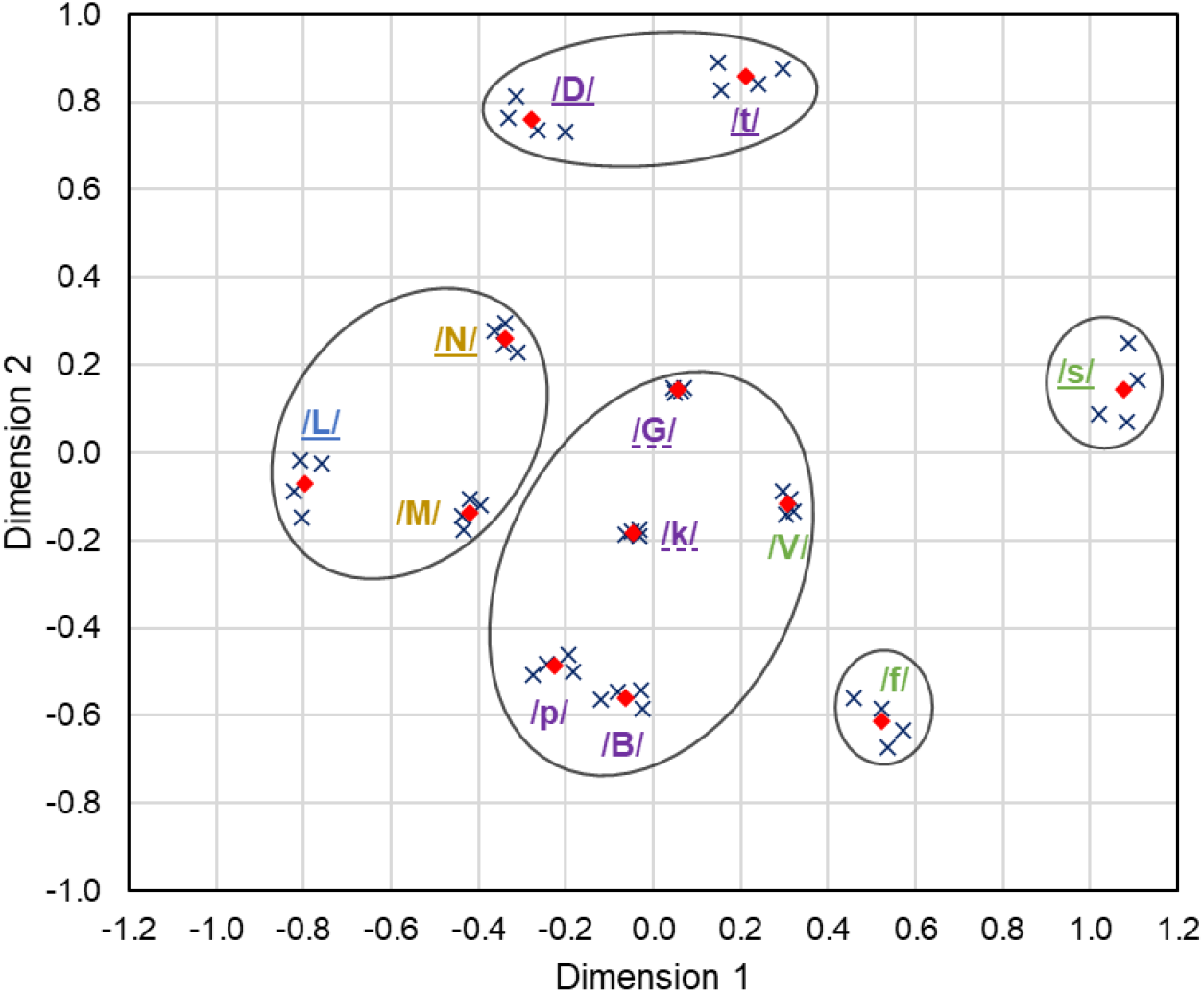
Shared perceptual map of the VCV-condition with the outer vowel /℧/ at +5 dB SNR. Blue crosses: Perceptual consonant coordinates of the different individuals. Red diamonds: Mean perceptual consonant coordinates of all individuals. Black circles: Consonant clusters according to the hierarchical cluster analysis.

**Supplementary Material N:**
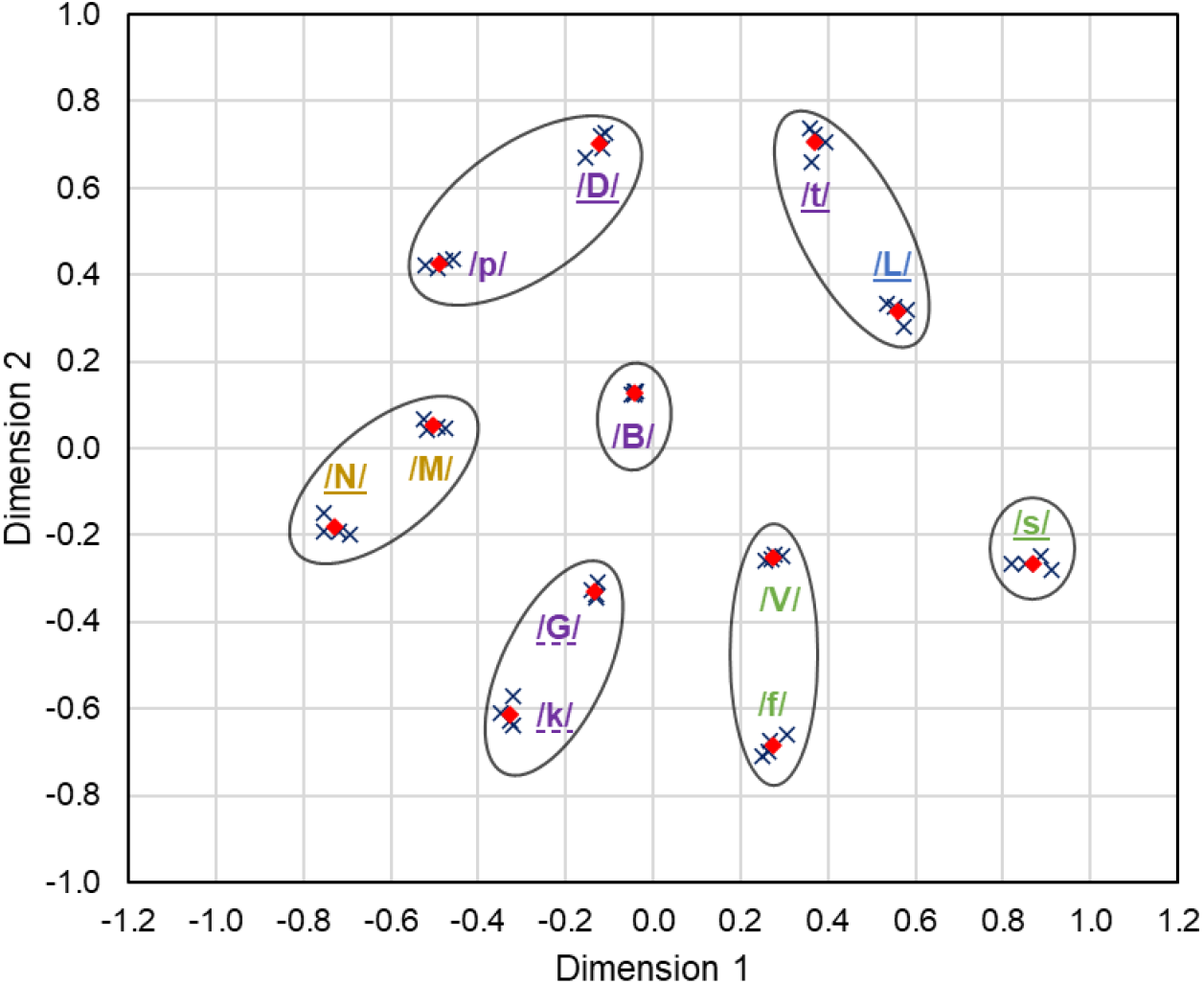
Shared perceptual map of the VCV-condition with the outer vowel /℧/ at +15 dB SNR. Blue crosses: Perceptual consonant coordinates of the different individuals. Red diamonds: Mean perceptual consonant coordinates of all individuals. Black circles: Consonant clusters according to the hierarchical cluster analysis.

## Footnotes

1 CVC – Consonant-vowel-consonant combination

2 VCV – Vowel-consonant-vowel combination

